# Comparison of beamformer implementations for MEG source localization

**DOI:** 10.1101/795799

**Authors:** Amit Jaiswal, Jukka Nenonen, Matti Stenroos, Alexandre Gramfort, Sarang S. Dalal, Britta U. Westner, Vladimir Litvak, John C. Mosher, Jan-Mathijs Schoffelen, Caroline Witton, Robert Oostenveld, Lauri Parkkonen

## Abstract

Beamformers are applied for estimating spatiotemporal characteristics of neuronal sources underlying measured MEG/EEG signals. Several MEG analysis toolboxes include an implementation of a linearly constrained minimum-variance (LCMV) beamformer. However, differences in implementations and in their results complicate the selection and application of beamformers and may hinder their wider adoption in research and clinical use. Additionally, combinations of different MEG sensor types (such as magnetometers and planar gradiometers) and application of preprocessing methods for interference suppression, such as signal space separation (SSS), can affect the results in different ways for different implementations. So far, a systematic evaluation of the different implementations has not been performed. Here, we compared the localization performance of the LCMV beamformer pipelines in four widely used open-source toolboxes (FieldTrip, SPM12, Brainstorm, and MNE-Python) using datasets both with and without SSS interference suppression.

We analyzed MEG data that were i) simulated, ii) recorded from a static and moving phantom, and iii) recorded from a healthy volunteer receiving auditory, visual, and somatosensory stimulation. We also investigated the effects of SSS and the combination of the magnetometer and gradiometer signals. We quantified how localization error and point-spread volume vary with SNR in all four toolboxes.

When applied carefully to MEG data with a typical SNR (3–15 dB), all four toolboxes localized the sources reliably; however, they differed in their sensitivity to preprocessing parameters. As expected, localizations were highly unreliable at very low SNR, but we found high localization error also at very high SNRs. We also found that the SNR improvement offered by SSS led to more accurate localization.

## 1. Introduction

MEG (magnetoencephalography) and EEG (electroencephalography) source imaging aims to identify the spatiotemporal characteristics of neural source currents based on the recorded signals, electromagnetic forward models and physiologically motivated assumptions about the source distribution. One well-known method for estimating a small number of focal sources is to model each of them as a current dipole with fixed location and fixed or changing orientation. The locations (optionally orientations) and time courses of the dipoles are then collectively estimated (Mosher et al., 1992; Hämäläinen et al., 1993). Such equivalent dipole models have been widely applied in basic research (see e.g. Salmelin, 2010) as well as in clinical practice (Bagic et al., 2011a; 2011b; Burgess et al., 2011). Distributed source imaging estimates source currents distribution across the whole source space, typically the cortical surface. Examples of linear methods for distributed source estimation are LORETA (low-resolution brain electromagnetic tomography; Pascual-Marqui et al., 1994) and MNE (minimum-norm estimation; Hämäläinen and Ilmoniemi, 1994). From estimated source distributions, one often computes noise-normalized estimates such as dSPM (dynamic statistical parametric mapping; Dale et al., 2000). Also, various non-linear distributed inverse methods have been proposed (Wipf et al., 2010; Gramfort et al., 2013b).

While dipole modeling and distributed source imaging estimate source distributions that reconstruct (the relevant part of) the measurement, beamforming takes an adaptive spatial-filtering approach, scanning independently each location in a predefined region of interest (ROI) within the source space without attempting to reconstruct the data. Beamforming can be done in time-or frequency domain; time-domain methods are typically based on the LCMV approach (Van Veen and Buckley, 1988; 1997; Spencer et al., 1992; Sekihara et al., 2006), and in frequency domain the DICS (Dynamic Imaging of Coherent Sources) (Gross et al., 2001) approach is popular.

The LCMV beamformer estimates the activity for a source at a given location (typically a point source) while simultaneously suppressing the contributions from all other sources and noise captured in the data covariance matrix. For evaluation of the spatial distribution of the estimated source activity, an image is formed by scanning a set of predefined possible source locations and computing the beamformer output (often power) at each location in the scanning space. When the scanning is done in a volume grid, the beamformer output is typically presented by superimposing it onto an anatomical MRI.

Beamformers have been popular in basic MEG research studies (e.g. Hillebrand and Barnes, 2005; Braca et al., 2011; Ishii et al., 2014; van Es and Schoffelen, 2019) as well as in clinical applications such as in localization of epileptic events (e.g. Van Klink et al., 2017; Youssofzadeh et al., 2018; Hall et al., 2018). Many variants of beamformers are implemented in several open-source toolboxes and commercial software for MEG/EEG analysis. Presently, based on citation counts, the most used open-source toolboxes for MEG data analysis are FieldTrip (Oostenveld et al., 2011), Brainstorm (Tadel et al., 2011), MNE-Python (Gramfort et al., 2013a) and DAiSS in SPM12 (Litvak et al., 2011). These four toolboxes have an implementation of an LCMV beamformer, based on the same theoretical framework (van Veen et al., 1997; Sekihara et al., 2006). Yet, it has been anecdotally reported that these toolboxes may yield different results for the same data. These differences may arise not only from the core of the beamformer implementation but also from the previous steps in the analysis pipeline, including data import, preprocessing, forward model computation, combination of data from different sensor types, covariance estimation, and regularization method. Beamforming results obtained from the same toolbox may also differ substantially depending on the applied preprocessing methods; for example, Signal Space Separation (SSS; Taulu and Kajola 2005) reduces the rank of the data, which could affect beamformer output unpredictably if not appropriately considered in the implementation.

In this study, we evaluated the LCMV beamformer pipelines in the four open-source toolboxes and investigated the reasons for possible inconsistencies, which hinder the wider adoption of beamformers to research and clinical use where accurate localization of sources is required, e.g., in pre-surgical evaluation. These issues motivated us to study the conditions in which these toolboxes succeed and fail to provide systematic results for the same data and to investigate the underlying reasons.

## 2. Materials and Methods

### 2.1. Datasets

To compare the beamformer implementations, we employed MEG data obtained from simulations, phantom measurements, and measurements of a healthy volunteer who received auditory, visual, and somatosensory stimuli. For all human data recordings, informed consent was obtained from all study subjects in agreement with the approval of the local ethics committee.

#### 2.1.1. MEG systems

All MEG recordings were performed in a magnetically shielded room with a 306-channel MEG system (either Elekta Neuromag® VectorView or TRIUX™; Megin Oy, Helsinki, Finland), which samples the magnetic field distribution by 510 coils at distinct locations above the scalp. The coils are configured into 306 independent channels arranged on 102 triple-sensor elements, each housing a magnetometer and two perpendicular planar gradiometers. The location of the phantom or subject’s head relative to the MEG sensor array was determined using four or five head position indicator (HPI) coils attached to the scalp. A Polhemus Fastrak® system (Colchester, VT, USA) was used for digitizing three anatomical landmarks (nasion, left and right preauricular points) to define the head coordinate system. Additionally, the centers of the HPI coils and a set of ∼50 additional points defining the scalp were also digitized. The head position in the MEG helmet was determined at the beginning of each measurement using the ‘single-shot’ HPI procedure, where the coils are activated briefly, and the coil positions are estimated from the measured signals. The location and orientation of the head with respect to the helmet can then be calculated since the coil locations were known both in the head and in the device coordinate systems. After this initial head position measurement, continuous tracking of head movements (cHPI) was engaged by keeping the HPI coils activated to track the movement continuously.

#### 2.1.2. Simulated MEG data

To obtain realistic MEG data with known sources, we superimposed simulated sensor signals based on forward modeling of dipolar sources onto measured spontaneous MEG data utilizing a special in-house simulation software. Structural MRI images, acquired from a healthy adult volunteer using a 3-tesla MRI scanner (Siemens Trio, Erlangen, Germany), were segmented using the MRI Segmentation Software of Megin Oy (Helsinki, Finland) and the surface enveloping the brain compartment was tessellated with triangles (5-mm side length). Using this mesh, a realistic single-shell volume conductor model was constructed using the Boundary Element Method (BEM; Hämäläinen and Sarvas, 1989) implemented in the Source modeling software of Megin Oy. We also segmented the cortical mantle with the FreeSurfer software (Dale et al., 1999; Fischl et al., 1999; Fischl, 2012) for deriving a realistic source space. By using the “ico4” subdivision in MNE-Python, we obtained a source space comprising 2560 dipoles (average spacing 6.2 mm) in each hemisphere (Fig. 1). Out of these, we selected 25 roughly uniformly distributed source locations in the left hemisphere for the simulations (Fig. 1). All these points were at least 7.5 mm inwards from the surface of the volume conductor model. We activated each of the 25 dipoles – one at a time – with a 10-Hz sinusoid of 200-ms duration (2 cycles). The dipoles were simulated at eight source 136 amplitudes: 10, 30, 80, 200, 300, 450, 600 and 800 nAm.

**Fig. 1.**
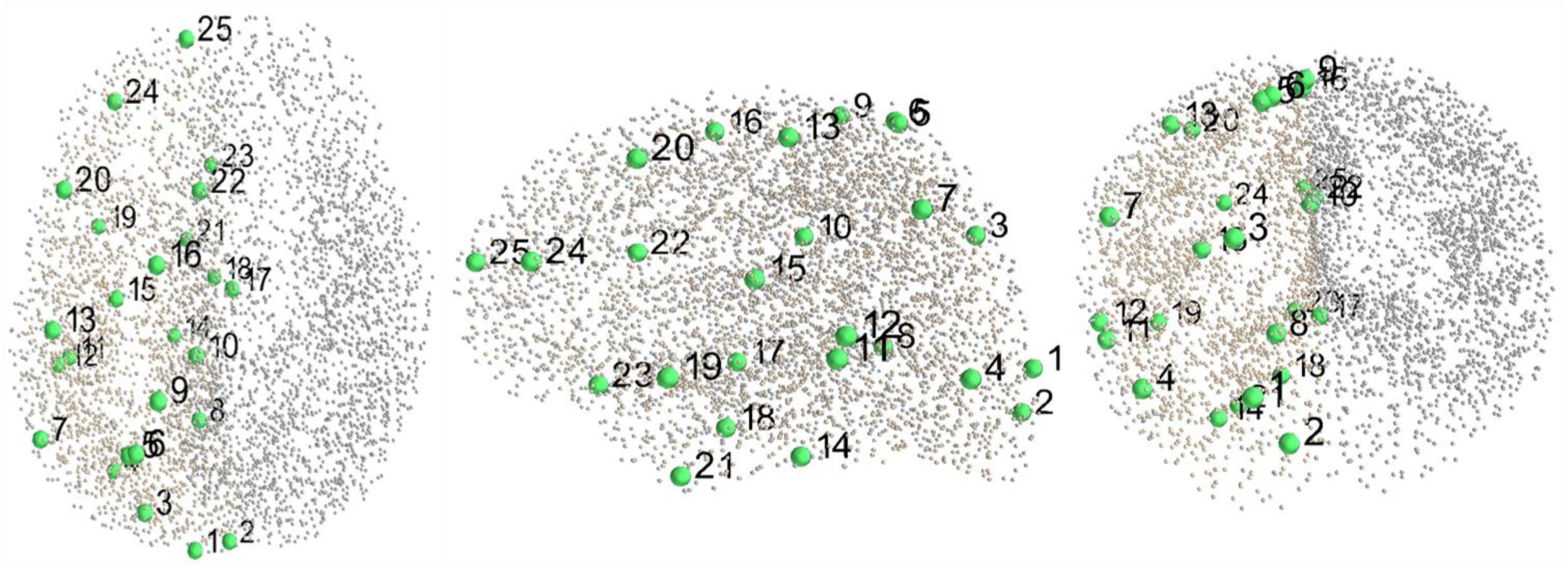
The 25 simulated dipolar sources (green dots) in the source space (grey dots).

A continuous resting-state MEG data with eyes open was recorded from the same volunteer who provided the anatomical data, using an Elekta Neuromag® MEG system (at BioMag Laboratory, Helsinki, Finland). The recording length was 2 minutes, the sampling rate was 1 kHz, and the acquisition frequency band was 0.1–330 Hz. This recording provided the head position for the simulations and defined their noise characteristics. MEG and MRI data were co-registered using the digitized head shape points and the outer skin surface in the segmented MRI.

The simulated sensor-level evoked fields were superimposed on the unprocessed resting-state recording with inter-trial-interval varying between 1000–1200 ms resulting in ∼110 trials (epochs) in each simulated dataset. The resting-state recording was used both as raw without preprocessing and after SSS interference suppression. Altogether, we obtained 400 simulated MEG datasets (25 source locations at 8 dipole amplitudes, all both with the raw and SSS-preprocessed real data). Fig. 2 illustrates the generation of simulated MEG data.

**Fig. 2.**
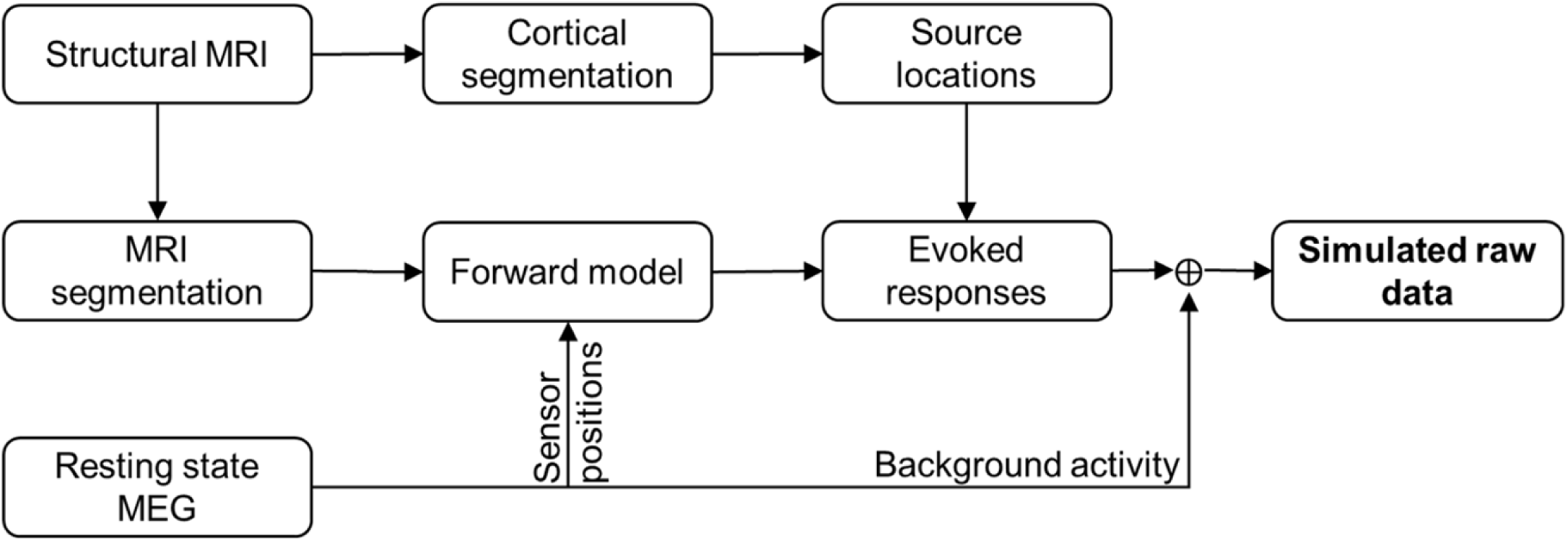
Simulating MEG data (detailed workflow in Suppl. Fig. 1).

#### 2.1.3. Phantom data

We used a commercial MEG phantom (Megin Oy, Helsinki, Finland) which contains 32 dipoles and 4 HPI coils at distinct fixed locations (see Fig 3a–c and Elekta Neuromag® TRIUX™ User’s Manual). The phantom is based on the triangle construction (Ilmoniemi et al., 1985): an isosceles triangular line current generates on its relatively very short side a magnetic field distribution equivalent to that of a tangential current dipole in a spherical conductor model, provided that the vertex of the triangle and the origin of the model of a conducting sphere coincide. The phantom data were recorded from 8 dipoles, excited one by one (see Elekta Neuromag® TRIUX™ User’s Manual), using a 306-channel TRIUX™ system (at Aston University, Birmingham, UK). The distance from the phantom origin was 64 mm for dipoles 5 and 9 (the shallowest), 54 mm for dipoles 6 and 10, 44 mm for dipoles 7 and 11, and 34 mm for dipoles 8 and 12 (the deepest; see Fig 3c). The phantom was first kept stationary inside the MEG helmet and continuous MEG data were recorded with 1-kHz sampling rate for three dipole amplitudes (20, 200 and 1000 nAm); one dipole at a time was excited with a 20-Hz sinusoidal current for 500 ms, followed by 500 ms of inactivity. The recordings were repeated with the 200-nAm dipole strength while moving the phantom continuously to mimic head movements inside the MEG helmet; see the movements in Fig. 3e and Suppl. Fig. 2 for all movement parameters.

**Fig. 3.**
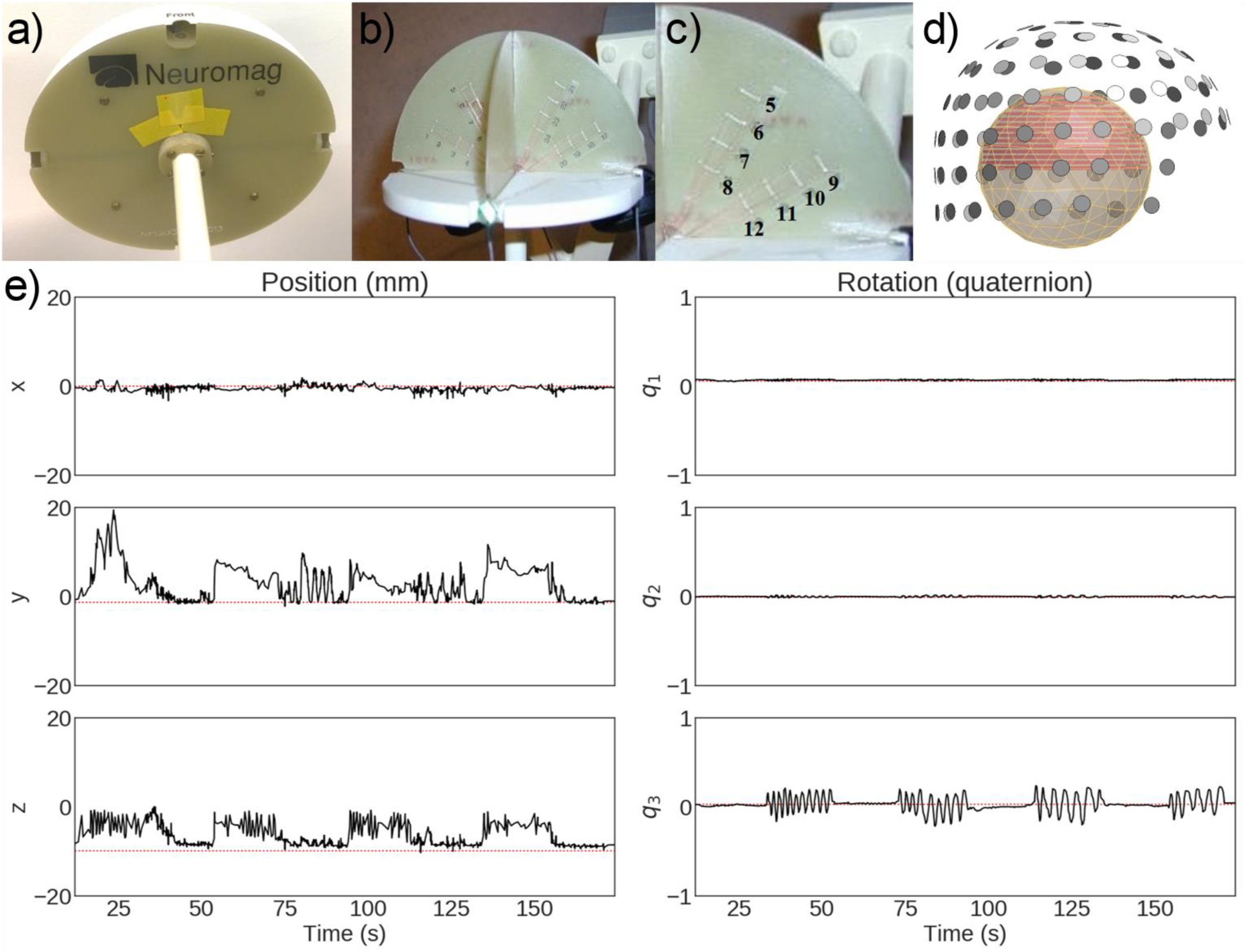
The dry phantom measured in this study. (a) Outer view, (b) cross-section, (c) positions of the employed dipole sources, (d) phantom position with respect to the MEG sensor helmet, and (e) position and rotation of the phantom during one of the moving-phantom measurements (Dipole 9 activated).

#### 2.1.4. Human MEG data

We recorded MEG evoked responses from the same volunteer whose MRI and spontaneous MEG data were utilized in the simulations. These human data were recorded using a 306-channel Elekta Neuromag® system (at BioMag Laboratory, Helsinki, Finland). During the MEG acquisition, the subject was receiving a random sequence of visual (a checkerboard pattern in one of the four quadrants of the visual field), somatosensory (electric stimulation of the median nerve at the left/right wrist at the motor threshold) and auditory (1-kHz 50-ms tone pips to the left/right ear) stimuli with an interstimulus interval of ∼500 ms. The Presentation software (Neurobehavioral Systems, Inc., Albany, CA, USA) was used to produce the stimuli.

### 2.2. Preprocessing

The datasets were analyzed in two ways: 1) omitting bad channels from the analysis, without applying SSS preprocessing, and 2) applying SSS-based preprocessing methods (SSS/tSSS) to reduce magnetic interference and perform movement compensation for moving phantom data. The SSS-based preprocessing and movement compensation were performed in MaxFilter™ software (version 2.2; Megin Oy, Helsinki, Finland). After that, the continuous data were bandpass filtered (passband indicated for each dataset later in the text) followed by the removing the dc. Then the data were epoched to trials around each stimulus. We applied an automatic trial rejection technique based on the maximum variance across all channels, rejecting trials that had variance higher than the 98^th^ percentile of the maximum or lower than the 2^nd^ percentile (see Suppl. Fig. 4). This method is available as an optional preprocessing step in FieldTrip, and the same implementation was applied in the other toolboxes. Below we describe the detailed preprocessing steps for all datasets.

#### 2.2.1. Simulated data

In each toolbox, the raw data with just bad channels removed or SSS-preprocessed continuous data were filtered using a zero-phase filter with a passband of 2–40 Hz. The filtered data were epoched into windows from –200 to +200 ms relative to the start of the source activity. The bad epochs were removed using the variance-based automatic trial rejection technique, resulting in ∼100 epochs. Then the noise and data covariance matrices were estimated from these epochs for the time windows of –200 to –20 ms and 20 to 200 ms, respectively.

#### 2.2.2. Phantom data

All 32 datasets (static: 3 dipole strengths and 8 dipole locations; moving: 1 dipole strength and 8 dipole locations) were analyzed both without and with SSS-preprocessing. We applied SSS on static phantom data to remove external interference. On moving-phantom data, combined temporal SSS and movement compensation (tSSS_mc) were applied for suppressing external and movement-related interference and for transforming the data from the continuously estimated positions into a static reference position (Taulu and Kajola 2005; Nenonen et al., 2012). Then in each toolbox the continuous data were filtered to 2–40 Hz using a zero-phase bandpass filter, and the filtered data were epoched from –500 to +500 ms with respect to stimulus triggers. Bad epochs were removed using the automated method based on maximum variance, yielding ∼100 epochs for each dataset. The noise and data covariance matrices were estimated in each toolbox for the time windows of – 500 to –50 ms and 50 to 500 ms, respectively.

#### 2.2.3. Human MEG data

Both the unprocessed raw data and the data preprocessed with tSSS were filtered to 1–95 Hz using a zero-phase bandpass filter in each toolbox. The trials with somatosensory stimuli (SEF) were epoched between –100 to –10 and 10 to 100 ms for estimating the noise and data covariances, respectively. The corresponding time windows for the auditory-stimulus trials (AEF) were –150 to – 20 and 20 to 150 ms, and for the visual stimulus trials (VEF) –200 to –50 and 50 to 200 ms, respectively. Trials contaminated by excessive eye blinks (EOG > 250 μV) or by excessive magnetic signals (MEG > 5000 fT or 3000 fT/cm) were removed with the variance-based automated trial removal technique. Before covariance computation, baseline correction by the time window before the stimulus was applied on each trial. The covariance matrices were estimated independently in each toolbox.

Since the actual source locations associated with the evoked fields are not precisely known, we defined reference locations using conventional dipole fitting in the Source Modeling Software of Megin Oy (Helsinki, Finland). A single equivalent dipole was used to represent SEF and VEF sources, and one dipole per hemisphere was used for AEF (see Suppl. Fig. 3). The dipole fitting was performed at the time point of the maximum RMS value across all planar gradiometer channels (global field power) of the average response amplitude.

#### 2.2.4. Forward model

For the beamformer scan of simulated data, we used the default or the most commonly used forward model of each toolbox: a single-compartment BEM model in MNE-Python, a single-shell corrected-sphere model (Nolte, 2003) in FieldTrip, a single-shell corrected sphere model (Nolte, 2003) through inverse normalization of template meshes (Mattout et al., 2007) in SPM12(DAiSS), and the overlapping-spheres (Huang et al., 1999) model in Brainstorm. For constructing models for these forward solutions, the segmentation of MRI images was performed in FreeSurfer for MNE-Python and Brainstorm while FieldTrip and SPM12 used the SPM segmentation procedure. A volumetric source space was represented by a rectangular grid with 5-mm resolution and 5-mm minimal distance from the head model surface. Forward solutions were computed separately in each toolbox using the head model, the volumetric grid sources, and sensor information from the MEG data. Since each toolbox prepares a head model using a different method, the shape of the head models may slightly differ from each other (see Fig. 4) which further may result in a shift between the positions of the scanning grid in these toolboxes.

**Fig. 4:**
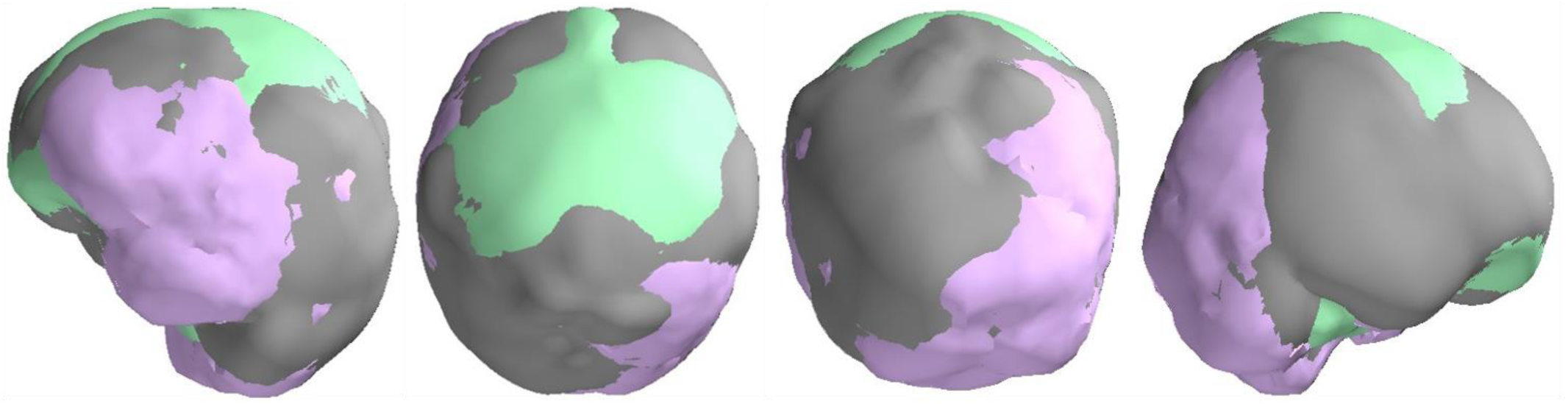
Correspondence between the head models used by MNE-Python (grey), FieldTrip (lavender) and SPM12 (mint). The Brainstorm head model is not included here as it uses overlapping spheres. The outermost surface (inner skull) across the toolboxes is rendered visible.

For phantom data, a homogeneous spherical volume conductor model was defined in each toolbox with the origin at the head coordinate system origin. An equidistant rectangular source-point grid with 5-mm resolution was placed inside the upper half of a sphere covering all 32 dipoles of the phantom; see Fig. 3d. Forward solutions for these grids were computed independently in each toolbox. For human MEG data, the head models and the source space were defined in the same way as for the beamformer scanning of the simulated data.

### 2.3. LCMV beamformer

The linearly constrained minimum-variance (LCMV) beamformer is a spatial filter that relates the magnetic field measured outside the head to the underlying neural activities using the covariance of measured signals and models of source activity and signal transfer between the source and the sensor (Spencer et al., 1992; van Veen et al. 1997; Robinson and Vrba, 1998). The spatial filter weights are computed for each location in the region of interest (ROI).

Let **x** be an *M* × 1 signal vector of MEG data measured with *M* sensors, and *N* is the number of grid points in the ROI with grid locations r_j_, (j = 1, …, *N*). Then the source **y**(*r*_*j*_) at any location *r*_*j*_ can be estimated as weighted combination of the measurement **x** as

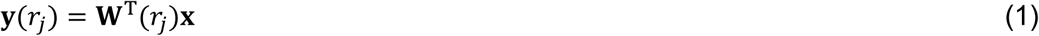

where the *M* × 3 matrix **W**(*r*_*j*_) is known as spatial filter for a source at location *r*_*j*_. This type of spatial filter provides a *vector type beamformer* by separately estimating the activity for three orthogonal source orientations, corresponding to the three columns of the matrix. According to Eqs 16–23 in van Veen et al. (1997), the spatial filter **W**(*r*_*j*_) for vector beamformer is defined as

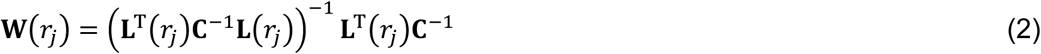

Here **L**(*r*_*j*_) is the *M* × 3 local leadfield matrix that defines the contribution of a dipole source at location *r*_*j*_ in the measured data **x**, and **C** is the covariance matrix computed from the measured data samples. To perform source localization using LCMV, the output variance (or output source power) Var(**y**(r_j_)) is estimated at each point in the source space (see Eq (24) in van Veen et al., 1997), resulting in

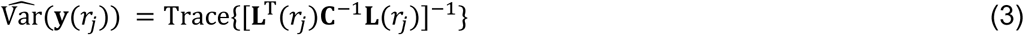

Usually, the measured signal is contaminated by non-uniformly distributed noise and therefore the estimated signal variance is often normalized with projected noise variance **C**_n_ calculated over some baseline data (noise). Such normalized estimate is called Neural Activity Index (NAI; van Veen et al., 1997) and can be expressed as

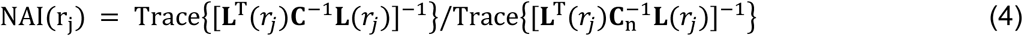

Scanning over all the locations in the region of interest in source space transforms the MEG data from a given measurement into an NAI map.

In contrast to a vector beamformer, a *scalar beamformer* (Sekihara and Scholz, 1996; Robinson and Vrba, 1998) uses constant source orientation that is either pre-fixed or optimized from the input data by finding the orientation that maximizes the output source power at each target location. Besides simplifying the output, the optimal-orientation scalar beamformer enhances the output SNR compared to the vector beamformer (Robinson and Vrba, 1998; Sekihara et al., 2004). The optimal orientation η_opt_(*r*_*j*_), for location *r*_*j*_ can be determined by generalized eigenvalue decomposition (Sekihara et al., 2004) using Rayleigh–Ritz formulation as

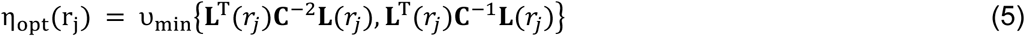

where *v*_min_ indicates the eigenvector corresponding to the smallest generalized eigenvalue of the matrices enclosed in Eq (5) curly braces. For further details, see Eq (4.44) and Section 13.3 in Sekihara and Nagarajan (2008).

Denoting 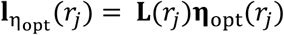 instead of **L**(*r*_*j*_), the weight matrix in Eq (2) becomes *M* × 1 weight vector ***w***(*r*_*j*_),

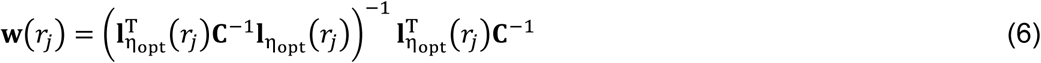

Using 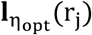 in Eq (4), we find the estimate (NAI) of a scalar LCMV beamformer as

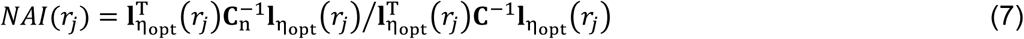

When the data covariance matrix is estimated from a sufficiently large number of samples and it has full rank, Eq (7) provides the maximum spatial resolution (Lin et al., 2008; Sekihara and Nagarajan, 2008). According to van Veen and colleagues (1997), the number of samples for covariance estimation should be at least three times the number of sensors. Thus, sometimes, the amount of available data may be insufficient to obtain a good estimate of the covariance matrices. In addition, pre-processing methods such as signal-space projection (SSP) or signal-space separation (SSS) reduce the rank of the data, which impacts the matrix inversions in Eq (7). These problems can be mitigated using Tikhonov regularization (Tikhonov, 1963) by replacing matrix **C**^−1^ by its regularized version (**C** + λ**I**)^−1^ in Eqs (2–7) where λ is called the regularization parameter.

All tested toolboxes set the λ with respect to the mean data variance, using ratio 0.05 as default:

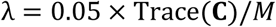

If the data are not full rank, also the noise covariance matrix **C**_n_ needs to be regularized.

### 2.4. Differences between the beamformer pipelines

Though all the four toolboxes evaluated here use the same theoretical framework of the LCMV beamformer, there are several implementation differences which might affect the exact outcome of a beamformer analysis pipeline. Many of these differences pertain to specific handling of the data prior to the estimation of the spatial filters, or to specific ways of (post)processing the beamformer output. Some of the toolbox-specific features reflect the characteristics of the MEG system around which the toolbox has evolved. Importantly, some of these differences are sensitive to input SNR, and they can lead to differences in the results. Table 1 lists the main characteristics and settings of the four toolboxes used in this study. We used the default settings of each toolbox (general practice) for steps before beamforming but set the actual beamforming steps as similar as possible across the toolboxes to be able to meaningfully compare the results.

**Table 1.**
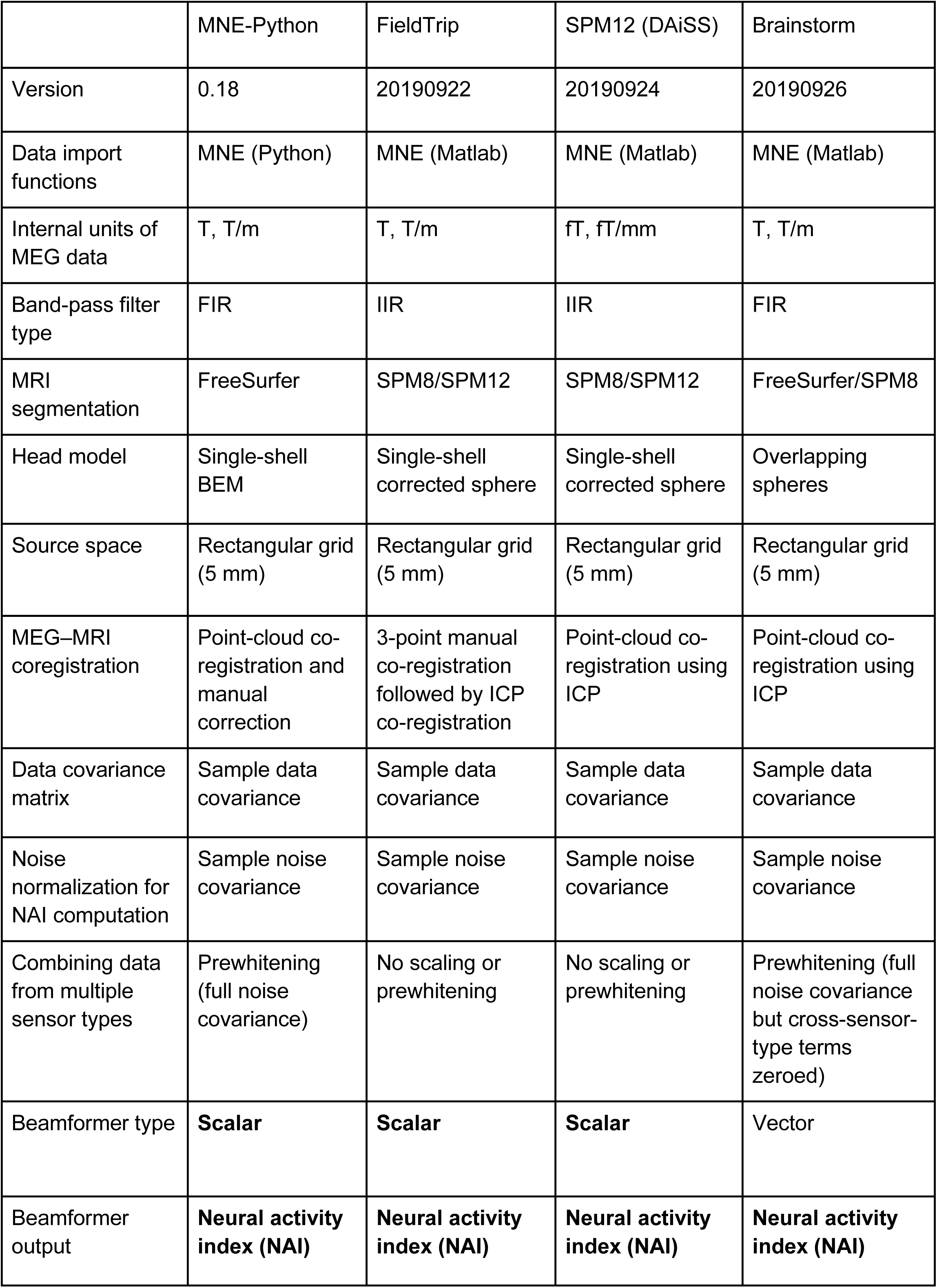
Characteristics of the four beamforming toolboxes. The non-default settings of each toolbox are shown in bold. The toolbox version is indicated either by the version number or by the download date (*yyyymmdd)* from GitHub.

|  | MNE-Python | FieldTrip | SPM12 (DAiSS) | Brainstorm |
| --- | --- | --- | --- | --- |
| Version | 0.18 | 20190922 | 20190924 | 20190926 |
| Data import functions | MNE (Python) | MNE (Matlab) | MNE (Matlab) | MNE (Matlab) |
| Internal units of MEG data | T, T/m | T, T/m | fT, fT/mm | T, T/m |
| Band-pass filter type | FIR | IIR | IIR | FIR |
| MRI segmentation | FreeSurfer | SPM8/SPM12 | SPM8/SPM12 | FreeSurfer/SPM8 |
| Head model | Single-shell BEM | Single-shell corrected sphere | Single-shell corrected sphere | Overlapping spheres |
| Source space | Rectangular grid (5 mm) | Rectangular grid (5 mm) | Rectangular grid (5 mm) | Rectangular grid (5 mm) |
| MEG–MRI coregistration | Point-cloud co-registration and manual correction | 3-point manual co-registration followed by ICP co-registration | Point-cloud co-registration using ICP | Point-cloud co-registration using ICP |
| Data covariance matrix | Sample data covariance | Sample data covariance | Sample data covariance | Sample data covariance |
| Noise normalization for NAI computation | Sample noise covariance | Sample noise covariance | Sample noise covariance | Sample noise covariance |
| Combining data from multiple sensor types | Prewhitening (full noise covariance) | No scaling or prewhitening | No scaling or prewhitening | Prewhitening (full noise covariance but cross-sensor-type terms zeroed) |
| Beamformer type | **Scalar** | **Scalar** | **Scalar** | Vector |
| Beamformer output | **Neural activity index (NAI)** | **Neural activity index (NAI)** | **Neural activity index (NAI)** | **Neural activity index (NAI)** |

All toolboxes import data using either Matlab or Python import functions of the MNE software (Gramfort et al., 2014) but represent the data internally either in T or fT (magnetometer) and T/m or fT/mm (gradiometer); see Suppl. Fig. 5. Default filtering approaches across toolboxes change the numeric values, so the linear correlation between the same channels across toolboxes deviates from the identity line; see Suppl. Fig. 6. The default head model is also different across toolboxes; see Section 2.2.4. The single-shell BEM and single-shell corrected sphere model (the “Nolte model”) are approximately as accurate but produce slightly different results (Stenroos et al., 2014).

For MEG–MRI co-registration, there are several approaches available across these toolboxes such as an interactive method using fiducial or/and digitization points defining the head surface, using automated point cloud registration methods e.g., the iterative closest point (ICP) algorithm. Despite using the same source-space specifications (rectangular grid with 5-mm resolution), differences in head models and/or co-registration methods change the forward model across toolboxes; see Fig. 4. Though there are several approaches to compute data and noise covariances across the four beamformer implementations, by default they all use the empirical/sample covariance. In contrast to other toolboxes, Brainstorm eliminates the cross-modality terms from the data and noise covariance matrices. Also, the regularization parameter *λ* is calculated and applied separately for gradiometers and magnetometers channel sets in Brainstorm.

The combination of two MEG sensor types in the MEGIN triple-sensor array causes additional processing differences in comparison to other MEG systems that employ only axial gradiometers or only magnetometers. Magnetometers and planar gradiometers have different dynamic ranges and measurement units, so their combination must be appropriately addressed in source analysis such as beamforming. For handling the two sensor types in the analysis, different strategies are used for bringing the channels into the same numerical range. MNE-Python and Brainstorm use pre-whitening (Engemann et al., 2015; Ilmoniemi and Sarvas, 2019) based on noise covariance while FieldTrip and SPM12 assume a single sensor type for all the MEG channels. This approach makes SPM12 to favor magnetometer data (with higher numeric values of magnetometer channels) and FieldTrip to favor gradiometer data (with higher numeric values of gradiometer channels). However, users of FieldTrip and SPM12 usually employ only one channel type of the triple-sensor array for beamforming (most commonly, the gradiometers). Due to the presence of two different sensor types in the MEGIN systems and the potential use of SSS methods, the eigenspectra of data from these systems can be idiosyncratic (see Suppl. Fig. 7) and differ from the single-sensor type MEG systems. Rank deficiency and related phenomena are potential sources of beamforming failures with data that have been cleaned with a method such as SSS.

Previous studies have shown that the scalar beamformer yields twofold higher output SNR compared to the vector-type beamformer, if the source orientation for the scalar beamformer has been optimized according to Eq 5 (Vrba J., 2000; Sekihara et al., 2004). Most of the beamformer analysis toolboxes have an implementation of optimal-orientation scalar beamformer. In this study, we used the scalar beamformer in MNE-Python, FieldTrip, and SPM12 but a vector-beamformer in Brainstorm since the orientation optimization was not available. To keep the output dimensionality the same across the toolboxes, we linearly summed the three-dimensional NAI values at each source location. The general workflow for analysis pipelines across toolboxes used in this study is illustrated in Suppl. Fig. 8.

### 2.5. Metrics used in comparison

In this study, a single focal source could be assumed to underlie the simulated/measured data. In such studies, accurate localization of the source is typically desired. We calculated two metrics for comparing the characteristics of the LCMV beamformer results from the four toolboxes: localization error, and point spread volume. We also analyzed their dependence on input signal-to-noise ratio.

#### Localization Error (LE)

True source locations were known for the simulated and phantom MEG data and served as reference locations in the comparisons. Since the exact source locations for the human subject MEG data were unknown, we applied the location of a single current dipole as a reference location (see Section 2.1.4 “Human MEG data”). The Source Modelling Software (Megin Oy, Helsinki, Finland) was used to fit a single dipole for each evoked-response category at the time point around the peak of the average response providing the maximum goodness-of-fit value. The beamformer localization error is computed as the Euclidean distance between the estimated and reference source locations.

#### Point-Spread Volume (PSV)

An ideal spatial filter should provide a unit response at the actual source location and zero response elsewhere. Due to noise and limited spatial selectivity, there is some filter leakage to the nearby locations, which spreads the estimated variance over a volume. The focality of the estimated source, also called focal width, depends on several factors such as the source strength, orientation, and distance from the sensors. PSV measures the focality of an estimate and is defined as the total volume occupied by the source activity above a threshold value; thus, a smaller PSV value indicates a more focal source estimate. We fixed the threshold to 50% of the highest NAI in all comparisons. In this study, the volume represented by a single source in any of the four source spaces (5-mm grid spacing) was 125 mm^3^.

#### Signal-to-Noise ratio (SNR)

Beamformer localization error depends on the input SNR, which varies – among other factors – as a function of source strength and distance of the source from the sensor array. Therefore, we evaluated beamformer localization errors and PSV as a function of the input SNR of the evoked field data.

We estimated the SNR for each evoked field MEG dataset in MNE-Python using the estimated noise covariance as follows: The data were whitened using the noise covariance and the effective number of sensors was then calculated as

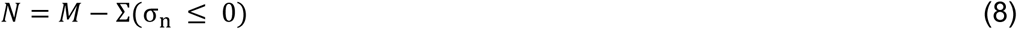

where *σ*_*n*_ are the eigenvalues of noise covariance matrix **C**_n_.

Then, the input SNR was calculated as:

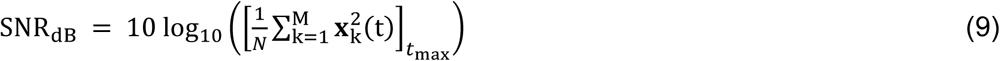

where x_k_(t) is the signal on the *k*^th^ sensor, *M* is the total number of sensors in the measurement, *t*_max_ is the time point at maximum amplitude of whitened data across all channels and *N* is the number of effective sensors defined in Eq (8). Since the same data were used in all toolboxes, we used the same input SNR value for all of them.

### 2.6. Data and code availability

The codes we wrote to conduct these analyses are publicly available under a repository https://zenodo.org/record/3471758 (DOI: 10.5281/zenodo.3471758). The datasets as well as the specific versions of the four toolboxes used in the study are available at https://zenodo.org/record/3233557 (DOI: 10.5281/zenodo.3233557).

## 3. Results

We computed the source localization error (LE) and the point spread volume (PSV) for each NAI estimate across all datasets from LCMV beamformer in all four toolboxes. We plotted the LE and PSV as a function of the input SNR computed according to Eq (9). To differentiate the localization among the implementations, we followed the following color convention: *MNE-Python: grey; FieldTrip: Lavender; SPM12 (DAiSS): Mint; and Brainstorm: coral*.

### 3.1. Simulated MEG data

Localization errors and PSV values were calculated for all simulated datasets and plotted against the corresponding input SNR. The SNR of all 200 simulated datasets ranged between 0.5 to 25 dB. Fig. 5a shows the plots between localization error and input SNR of each simulated dataset. The polynomial regressions of the maximum localization errors across LCMV implementations show the variation of localization errors over the range of input SNRs. The localization error goes high for all toolboxes for very low SNR (< 3 dB) signals (e.g. 20-nAm or deep sources). The localization error within the input SNR range 3–12 dB is stable and mostly within 15 mm, and SSS preprocessing widens this SNR range of stable performance to 3–15 dB. Unexpectedly, we also found high localization error at high SNR (> 15 dB) for the toolboxes other than SPM12 (DAiSS). Fig. 5b plots PSV values against input SNR for raw and SSS-preprocessed simulated data. The polynomial regression plots fit a nonlinear relationship between the input SNR and the corresponding maximal PSVs across the four LCMV implementations. The regression plots in Fig. 5b agree with the corresponding plots in Fig. 5a, i.e., lower PSV values (higher spatial resolution) for the SNR range with smaller localization errors and vice-versa, for all toolboxes. The low SNR signals (usually, weak or deep sources) shows high PSV values in Fig. 5b which also indicates improved spatial resolution after SSS preprocessing. Fig. 5a–b shows that none of the four toolboxes provides accurate localization for all SNR values and that the spatial resolution of LCMV is dependent on input SNR. SPM12 (DAiSS) shows lower localization errors and PSV values at very high SNR too.

**Fig. 5.**
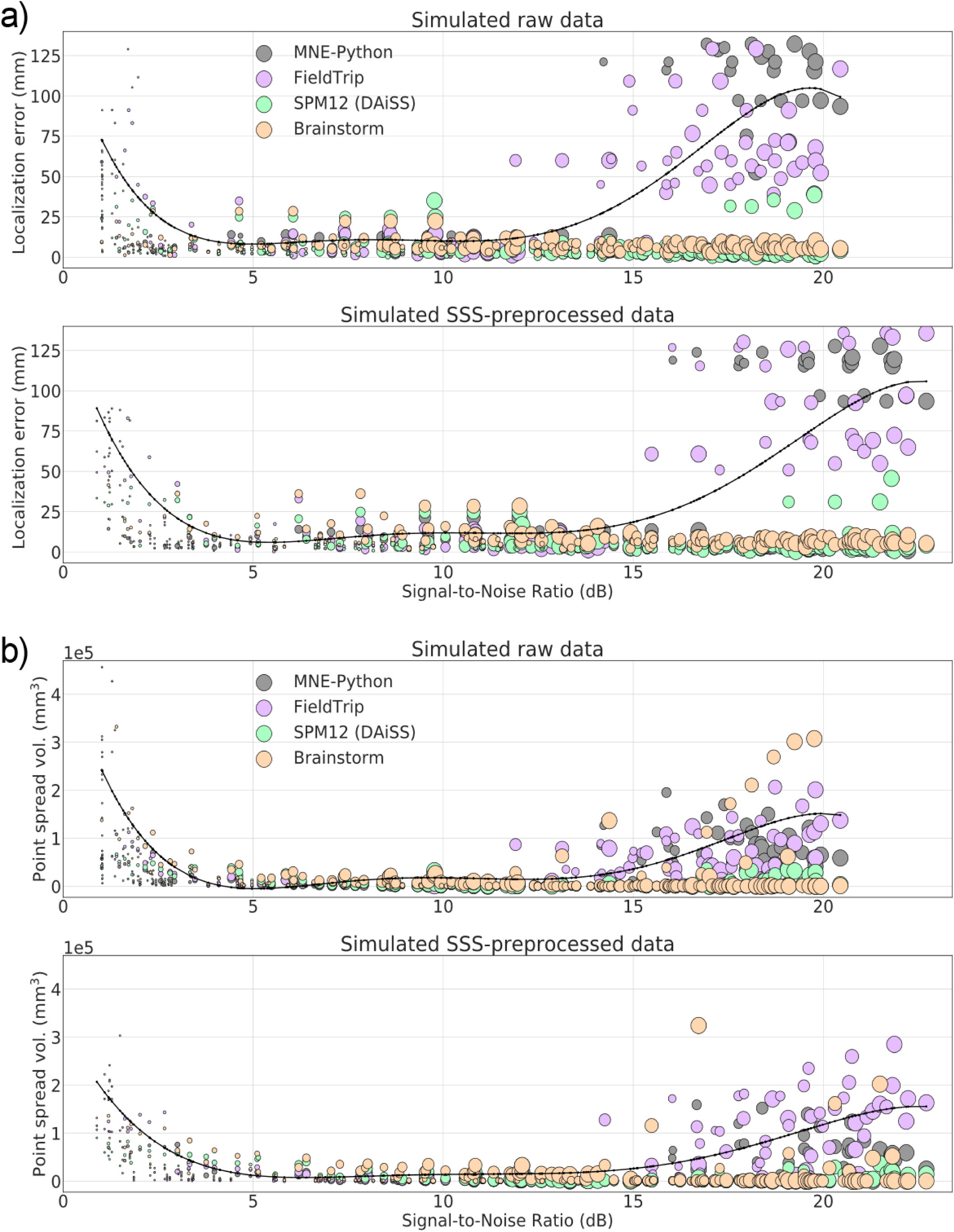
Localization error (a) and point-spread volume (b) as a function of input SNR for raw and SSS-preprocessed simulated datasets. The markers size indicates the true dipole amplitude. The curves (black) indicate the polynomial regression of the maximal value across the four LCMV implementations.

### 3.2. Static and moving phantom MEG data

In the case of phantom data, the background noise is very low and there is a single source underneath a measurement. Also, the phantom analysis uses a homogeneous sphere model that does not introduce any forward model inaccuracy, except the possible co-registration error. All four toolboxes show high localization accuracy and high resolution for phantom data, if the input SNR is not very low. Corresponding results for the static phantom data are presented in Fig. 6a–b. Fig. 6a indicates the localization error clear dependency on SNR. The nonlinear regression plots fitted over maximum localization errors indicate high localization errors at very low SNR raw data sets. The high error is because of some unfiltered artifacts in raw data which was removed by SSS. After SSS, the beamformer shows localization error under ∼5 mm for all the datasets. Fig. 6b shows the beamforming resolution in terms of PSV. The regression plots fitted over maximum PSV values show a high spatial resolution for the data with SNR > 5 dB.

**Fig. 6.**
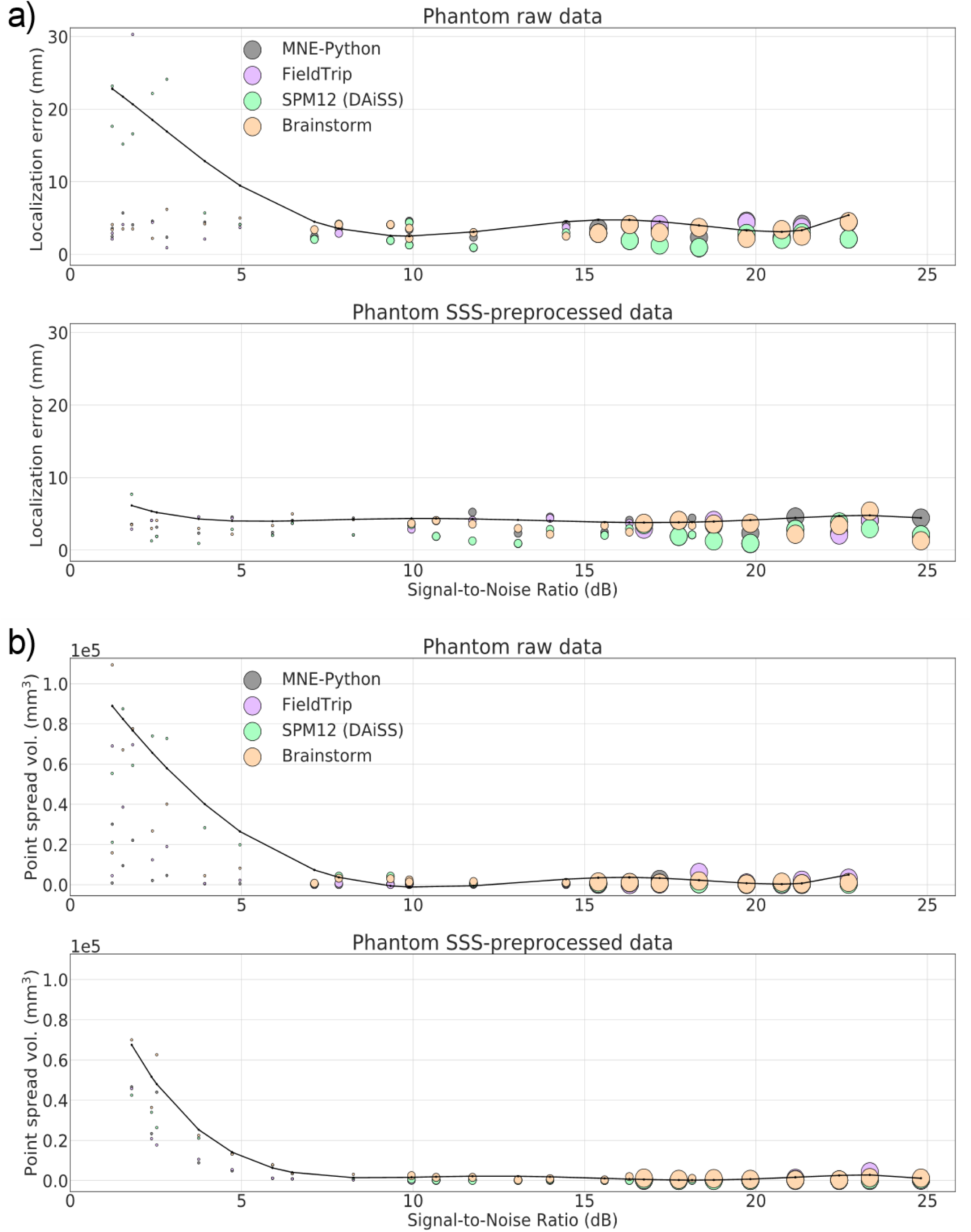
Localization error (a) and point-spread volume (b) as a function of input SNR for phantom data recording in a stable position. The markers size indicates the true dipole amplitude. The curves (black) indicate the polynomial regression of the across the four LCMV implementations.

In the cases of moving phantom, Fig. 7a shows high localization errors with unprocessed raw data because of disturbances caused by the movement. The dipole excitation amplitude was 200 nAm, which is enough to provide a good SNR. The most superficial dipoles (Dipoles 5 and 9 in Fig. 3c) possess higher SNR but also higher localization error since they get more significant angular displacement during movement. Because of differences in implementations and preprocessing parameters listed in Section 2.4, apparent differences among the estimated localization error can be seen. Overall, MNE-Python shows the lowest while SPM12 (DAiSS) shows the highest localization error with the phantom data with movement artifact. After applying for spatiotemporal tSSS and movement compensation, the improved SNR provided significantly better localization accuracies. Fig. 7b shows the PSV for moving phantom data for raw and processed data. The regression plots indicate improvement in SNR and spatial resolution after tSSS with movement compensation.

**Fig. 7.**
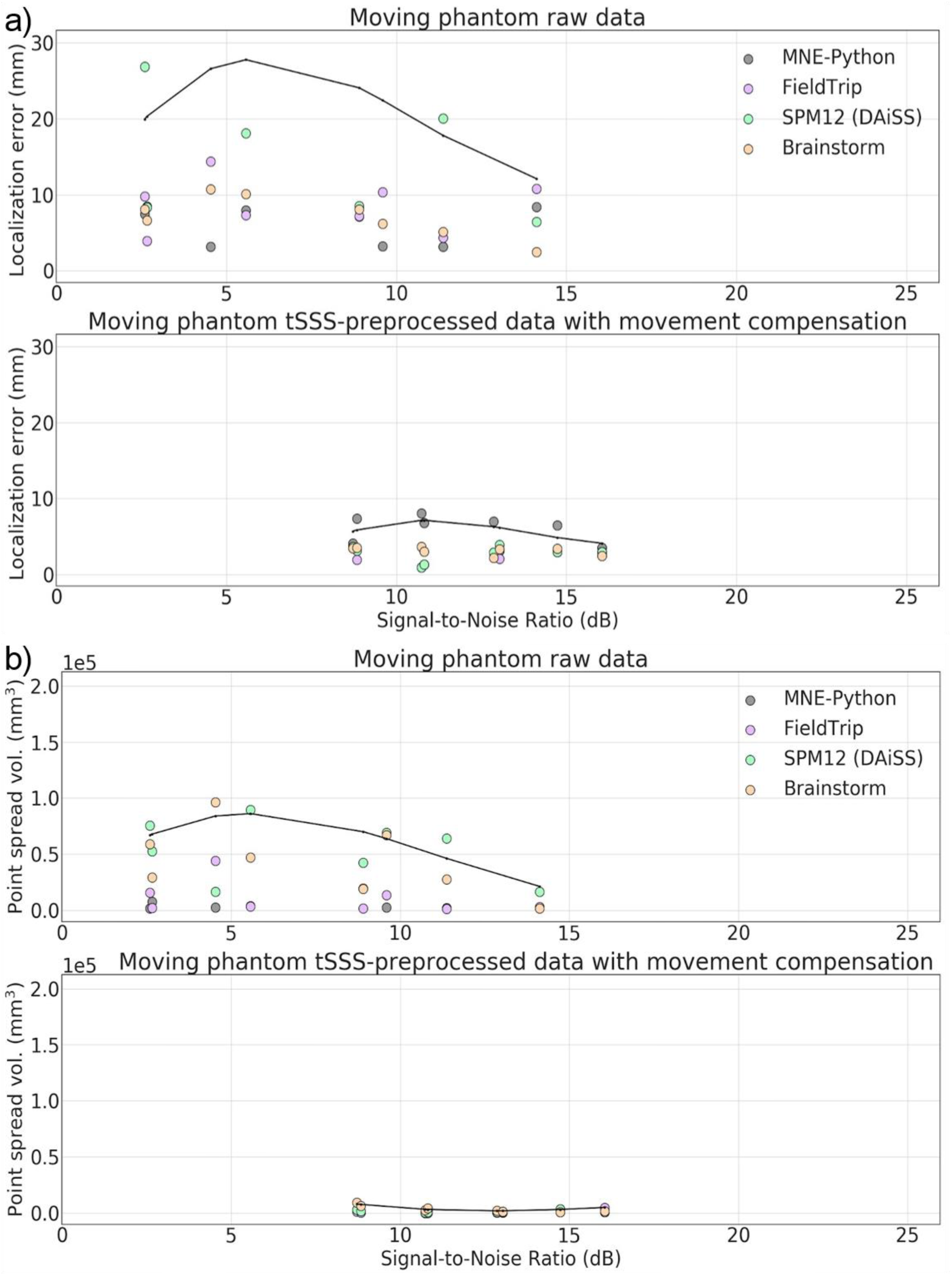
Localization error (a) and point-spread volume (b) as a function of input SNR for data from the moving phantom. The curves (black) indicate the polynomial regression of the maximum value across the four LCMV implementations.

### 3.3. Human subject MEG data

Since the correct source locations for the human evoked field datasets are unknown, we plotted the localization difference across the four LCMV implementations for each data. These localization differences were the Cartesian distance between an LCMV-estimated location and the corresponding reference dipole location as explained in Section 2.1.4. Fig. 8a shows the plots for the localization differences against the input SNRs computed using Eq (9) for four visual, two auditory and two somatosensory evoked-field datasets. The localization differences for both unprocessed raw and SSS preprocessed data are mostly under 20 mm in each toolbox. The higher differences compared to the phantom and simulated dataset could be because of two reasons. First, the recording might have been comprised by some head movement, which could not be corrected because of the lack of continuous HPI. Second, the reference dipole location may not represent the very same source as estimated by the LCMV beamformer. In contrast to dipole fitting, beamforming utilizes data from the full covariance window, so some difference between the estimated localizations is to be expected. For all SSS-preprocessed evoked field datasets, Fig. 8b shows the estimated locations across the four LCMV implementation and the corresponding reference dipole locations. For simplifying the visualization, all estimated locations in a stimulus category are projected onto a single axial slice. All localizations seem to be in the correct anatomical regions, except the estimated location from right-ear auditory responses by MNE-Python after SSS-preprocessing (Fig. 8b; red circle). After de-selecting the channels close to the right auditory cortex, the MNE-Python-estimated source location was correctly in the left cortex (Fig. 8b; green circle). The regression plots fitted over the maxima of the localization differences across the LCMV implementations show the improvement in input SNR and also localization improvement in some cases. Fig. 9 in Supplementary material shows the PSV values as a function of the input SNR for the evoked-field datasets, demonstrating the spatial resolution of beamforming.

**Fig. 8.**
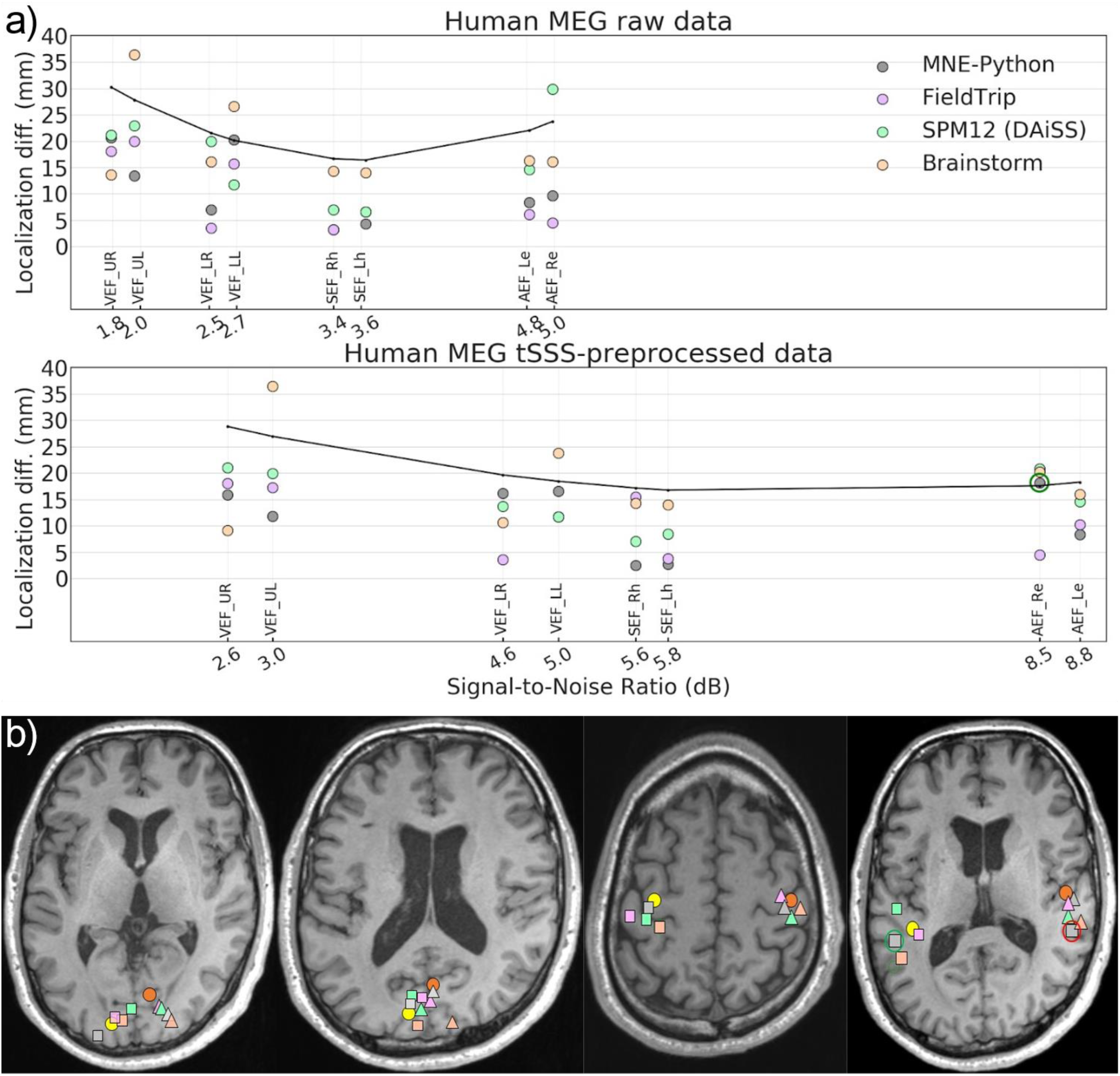
Source estimates of human MEG data. (a) Localization difference from the reference dipole location for raw and tSSS-preprocessed data. (b) Peaks of the beamformer source estimate of tSSS-processed data. From left to right: visual stimuli presented to left (triangle) and right (square) upper and lower quadrant of the visual field (the two axial slices showing all sources); somatosensory stimuli to left (triangle) and right (square) wrist; auditory stimuli to the left (triangle) and right (square) ear. Reference dipole locations (yellow and orange circles).

## 4. Discussion

The localization accuracy and beamformer resolution as a function of the input SNR were investigated and compared across the LCMV implementations in the four tested toolboxes. In the absence of background noise, the phantom data showed high localization accuracy and high spatial resolution if the input SNR >∼5 dB. All implementations also showed high localization accuracy for data recording from a moving phantom after compensating the movement and applying tSSS. For the simulated datasets with realistic background noise, the regression curve fitted over the maximum localization error across the LCMV implementations indicates that the reliability of localization accuracy in these implementations depends on the SNR of input data and these implementations localize a single source reliably within the SNR range of ∼3–15 dB. Small differences among the estimated source locations across the implementations even in this SNR range are caused by differing processing steps in defining the head model, spatial filter and performing the beamformer scan. For the human subject evoked-field MEG data, all implementations localize sources within about 20 mm.

Our results indicate that with the default parameter settings, none of the four implementations works universally reliable for all datasets and input SNR values. In the case of low SNR (typically less than 3 dB), the lower contrast between data and noise covariance may cause the beamformer scan to provide a flat peak in the output and so the localization error goes high. Unexpectedly, we found high localization error for high SNR signal and significant differences between the toolboxes. The regression curves fitted over averaged maximum PSV across all toolboxes showed higher values for low- and high-SNR simulated data. As expected, reliable localization provides higher spatial resolution across the implementations and vice-versa (Fig. 5 and 6). The lower spatial resolution (higher PSV) for the signal with low SNR also agrees with previous studies (Lin et al., 2008; Hillebrand and Barnes, 2003). We further discuss here the significant steps of the beamformer pipelines, which affect the localization accuracy and introduce discrepancies among the implementations.

### 4.1 Preprocessing with SSS

Due to the spatial-filter nature of the beamformer, it can reject external interference and therefore SSS-based pre-processing may have little effect on the results. Thus, although the SNR increases as a result of applying SSS, the localization accuracy does not necessarily improve, which is evident in the localization of the evoked responses (Fig. 8).

However, undetected artifacts, such as a large-amplitude signal jump in a single sensor, may in SSS processing spread to neighboring channels and subsequently reduce data quality. Therefore, channels with distinct artifacts should be noted and marked as bad prior to beamforming of unprocessed data or before applying SSS operations. In addition, trials with large artifacts should be removed based on an amplitude thresholding or by other means. Furthermore, SSS processing of extremely weak signals (SNR < ∼2 dB) may not improve the SNR for producing smaller localization errors and PSV values. Hence the data quality should be carefully inspected before and after applying preprocessing methods such as SSS, and channels or trials with low-quality data (or lower contrast) should be omitted from the covariance estimation.

### 4.2. Effect of filtering and artifact-removal methods

All four toolboxes we tested employ either a MATLAB or Python implementation of the same MNE routines (Gramfort et al. 2014) for reading FIFF data files and thus have internally the exact same data at the very first stage (see Suppl. Fig. 6). The data import either keeps the data in SI-units (T for magnetometers and T/m for gradiometers) or rescales the data (fT and fT/mm) before further processing. The actual pre-processing steps in the pipeline may contribute to differences in the results. The filtering step is performed to remove frequency components of no interest, such as slow drifts, from the data. By default, FieldTrip and SPM use an IIR (Butterworth) filter, and MNE-Python uses FIR filters. The power spectra of these filters’ output signals show notable differences and the output data from these two filters are not identical. Significant variations can be found between MNE-Python-filtered and FieldTrip/SPM-filtered data. Although SPM and FieldTrip use the same filter implementation, the filtering results are not identical because of numeric differences caused by different channel scaling (Suppl. Fig 6). These differences affect the estimated covariance matrices, which are a crucial ingredient for the spatial-filter computation and finally may contribute to differences in beamforming results.

### 4.3. Effect of SNR on localization accuracy

We reduced the impact of the unknown source depth and strength to a well-defined metrics in terms of the SNR. We observed that the localization accuracy is poor for very low SNR values, i.e. below 3 dB. The weaker, as well as the deeper sources, project less power on to the sensor array and thus show lower SNR; see Eq (9). On the other hand, the LCMV beamformer may also fail to localize accurately sources that produce very high SNR values, likely because the data covariance matrix is over-fitted, or the scanning grid is too sparse with respect to the point spread of the beamformer output. In this case the output is too focal and a small error in forward solution, introduced for example by inaccurate coregistration, may lead to missing the true focal source and obtaining nearly equal power estimates at many source locations, increasing the chance of mislocalization. Usually, such high levels of SNR do not occur in typical human MEG experiments, however, the strength of equivalent current dipoles (ECD) for interictal epileptiform discharges (IIEDs) typically ranges between 50 and 500 nAm (Bagic et al., 2011a).

All four beamformer pipelines provided very similar results when the SNR is in the “suitable range” of about ∼3–15 dB. Unsatisfactory performance is typically due to the data; either the SNR is extremely low, or there are some uncorrected artifacts in the data. The results of the phantom data showed that all toolboxes provide equally good results if there are no uncorrected large artifacts in the data and if the SNR is not extremely small or large.

### 4.4. Effect of the head model

Forward modelling requires MEG–MRI co-registration, segmentation of the head MRI and leadfield computation for the source space. The four beamformer implementations use different approaches, or similar approaches but with different parameters, which yields slightly different forward models. From Eqs (2–7), it is evident that beamformers are quite sensitive to the forward model. Hillebrand and Barnes (2003) showed that the spatial resolution and the localization accuracy of a beamformer improve with accuracy of the forward model. Dalal and colleagues (2014) reported that co-registration errors contribute greatly to EEG localization inaccuracy, likely due to their ultimate impact on head-model quality. Chella and colleagues (2019) presented the dependency of beamformer-based functional connectivity estimates on MEG-MRI co-registration accuracy.

The increasing inter-toolbox localization differences towards very low and very high input SNR is also subject to the differences between the head models. Fig. 4 shows the three overlapped head models prepared from the same MRI where a slight misalignment among head models can be easily seen. This misalignment also affects source space. These differences in head models and thus in forward solutions will contribute to differences in beamforming results across the toolboxes.

### 4.5. Covariance matrix

The data covariance matrix is a key component of the adaptive spatial filter in LCMV beamforming, and any error in covariance estimation can cause an error in source estimation. We used 5% of the mean variance of all sensors to regularize data covariance for making its inversion stable in FieldTrip, SPM12 and MNE-Python. Brainstorm uses a slightly different approach and applies regularization with 5% of mean variance of gradiometer and magnetometer channel sets separately and eliminate cross-sensor-type entries from the covariance matrices. As SSS preprocessing reduces the rank of the data, usually retaining at most 75 non-zero eigenvalues, the trace of the covariance matrix decreases strongly. At very high SNRs (> 15 dB), overfitting of the covariance matrix becomes more prominent; the condition number (ratio of the largest and the smallest eigenvalues) of the covariance matrix becomes very high even after the default regularization, which can deteriorate the quality of source estimates unless the covariance is appropriately regularized. Therefore, the seemingly same 5% regularization can have very different effects before and after SSS; see Suppl. Fig. 7. Thus, the commonly used way of specifying the regularization level might not be appropriate to produce a good and stable covariance model at high SNR, and this could be one of the explanations for the anecdotally reported detrimental effects of SSS on beamforming results.

## 5. Conclusion

We conclude that with the current versions of LCMV beamformer implementations in the four open-source toolboxes — FieldTrip, SPM12(DAiSS), Brainstorm, and MNE-Python — the localization accuracy is acceptable (within ∼10 mm for a true point source) for most purposes when the input SNR is 3–15 dB. Lower or higher SNR may compromise the localization accuracy and spatial resolution. To extend this useable range, a properly defined scaling strategy such as pre-whitening should be implemented across the toolboxes. The default regularization is often inadequate and may yield suboptimal results. Therefore, a data-driven approach for regularization should be adopted to alleviate problems with low- and high-SNR cases. Our further work will be focusing on optimizing regularization using a more data-driven approach.

## Acknowledgment

This study has been supported by the European Union H2020 MSCA-ITN-2014-ETN program, Advancing brain research in Children’s developmental neurocognitive disorders project (ChildBrain #641652). SSD and BUW have been supported by an ERC Starting Grant (#640448).

