## Supplemental for "Comparison of beamformer implementations for MEG source localization"

[1]Megin Oy, Helsinki, Finland.

[2]Department of Neuroscience and Biomedical Engineering, Aalto University School of Science, Espoo, Finland

[3]Inria, CEA/Neurospin, Universite Paris-Saclay, Paris, France

[4]Center of Functionally Integrative Neuroscience, Aarhus University, Denmark

[5]The Wellcome Centre for Human Neuroimaging, UCL Queen Square Institute of Neurology, London, UK

[6]Department of Neurology, University of Texas Health Science Center at Houston, Houston, Texas, USA

[7]Donders Institute for Brain, Cognition and Behaviour, Radboud University, Nijmegen, The Netherlands

[8]NatMEG, Karolinska Institutet, Stockholm, Sweden

[9]Aston Brain Centre, School of Life and Health Sciences, Aston University, Birmingham, UK

---

The data as well as the used versions of the four open-source beamforming toolboxes — FieldTrip, SPM12(DAiSS), Brainstorm, and MNE-Python are publicly available at repository https://zenodo.org/record/3233557 (DOI: 10.5281/zenodo.3233557). The availability of simulated data are limited to 50 (randomly selected) out of the total of 400 because of repository space constraint, the rest of the simulated data would be available on demand. Here we add all the supplementary information to provide more detailed information at various stages in our study.

### 1. Simulated data generation

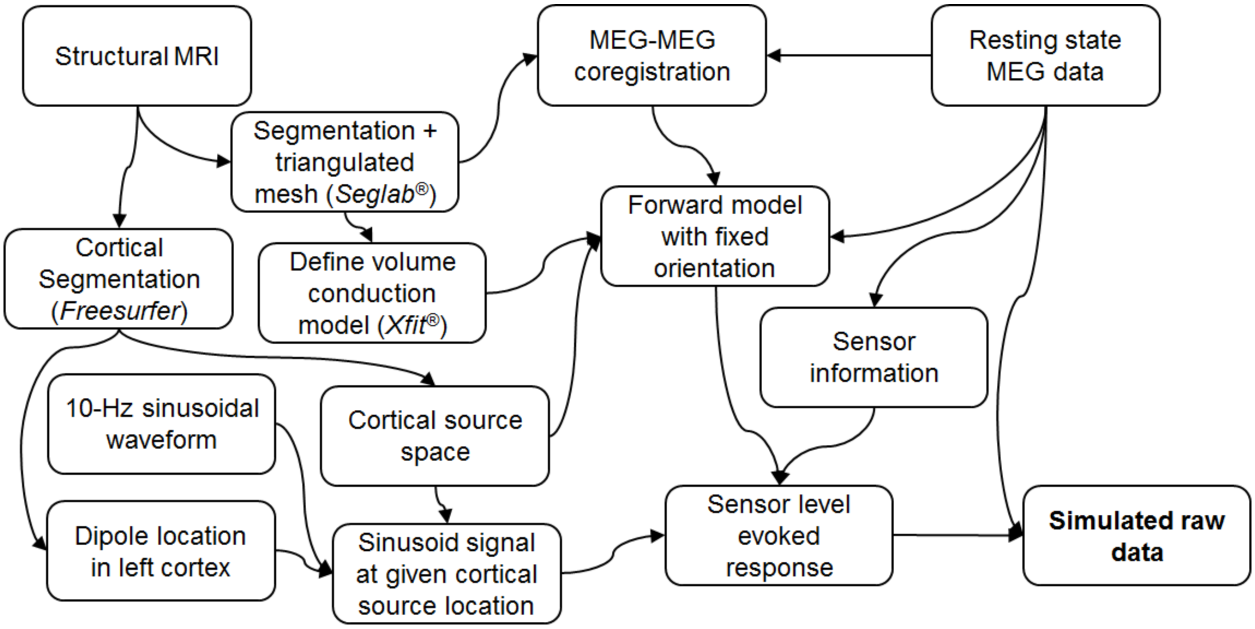


**Fig. 1.** Detailed workflow for simulating MEG data at 25 source locations

### 2. Phantom movement inside the MEG helmet

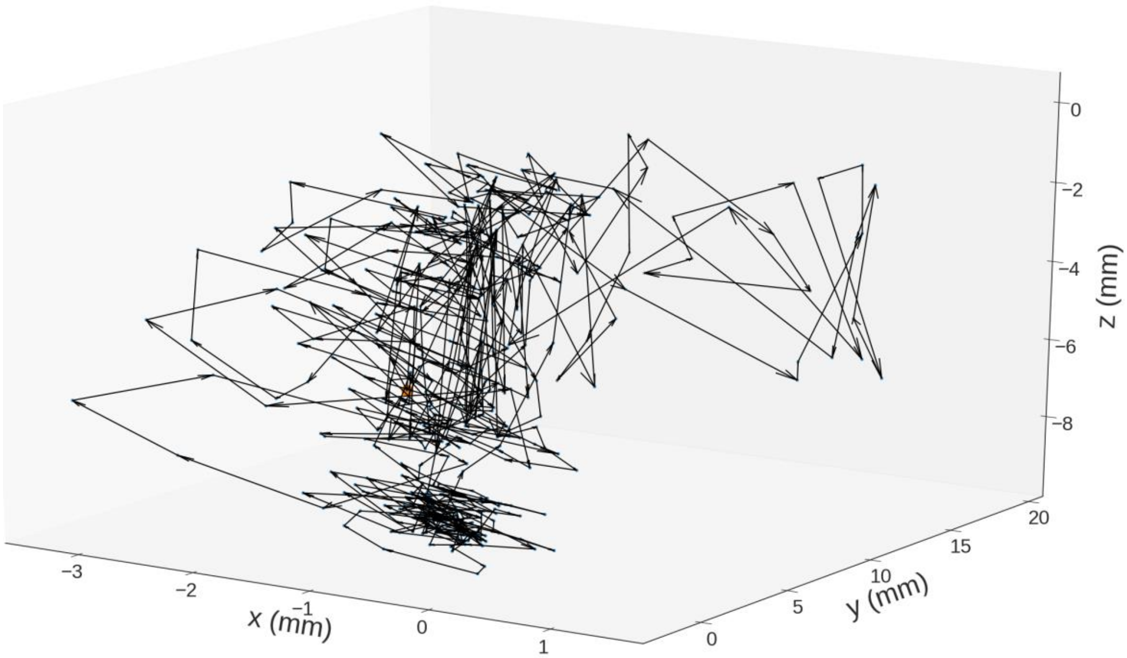


**Fig. 2.** 6D movements of phantom inside MEG helmet while recording moving phantom data (where dipole 9 was stimulated at 200 nAm).

**3. Dipole localization for multimodal data using Xfit(R)**

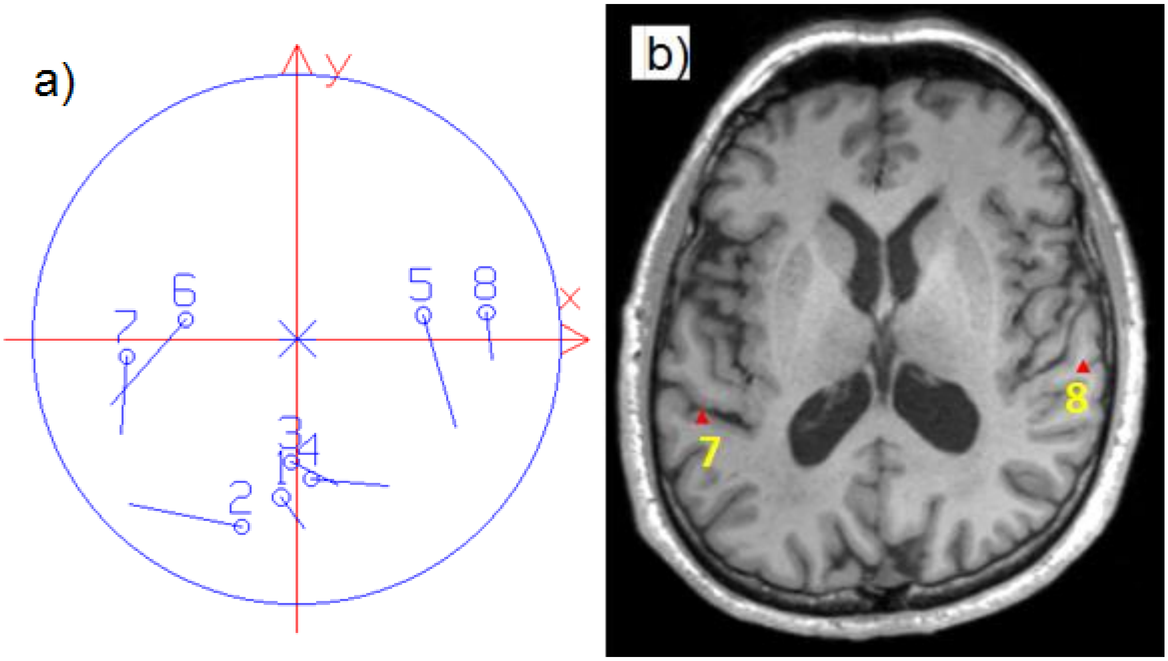

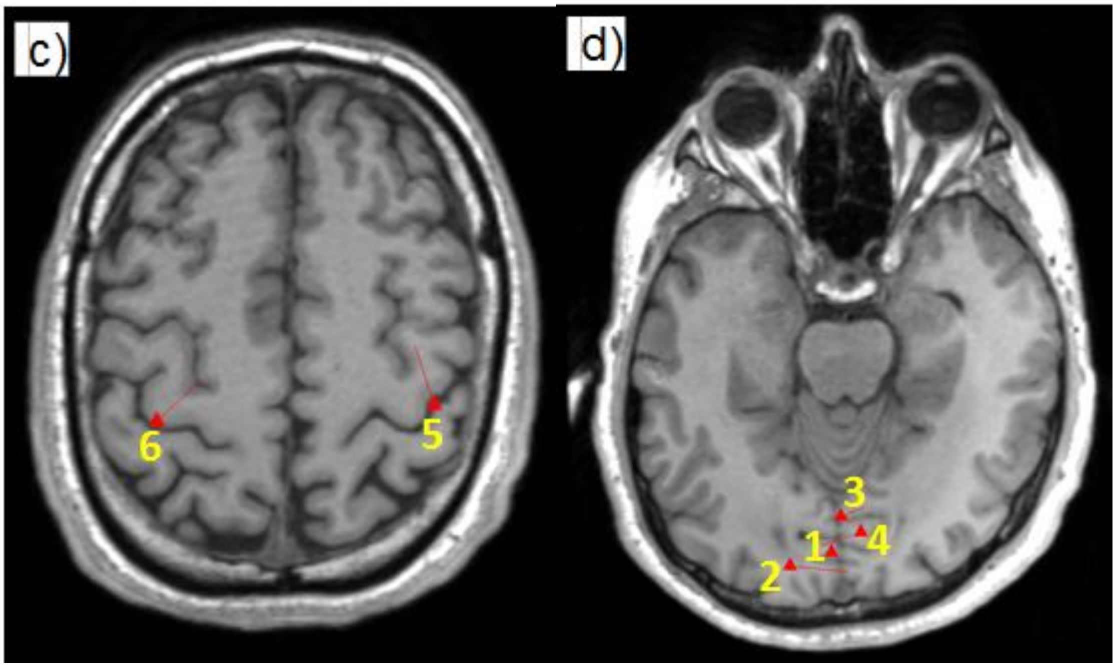


**Fig. 3.** The estimated dipoles for the multimodal data set. (a) All the eight dipoles projected on a common x-y plane; (b) dipoles 7 for auditory left ear stimulus and 8 for auditory right ear stimulus; (c) dipoles 6 for somatosensory stimuli in the left hand and 8 for the right hand; (d) dipoles 1–4 are for visual stimuli presented at upper-right, lower-right, upper-left and lower left quadrant of the stimulus screen respectively.

### 4. Covariance-based epoch rejection

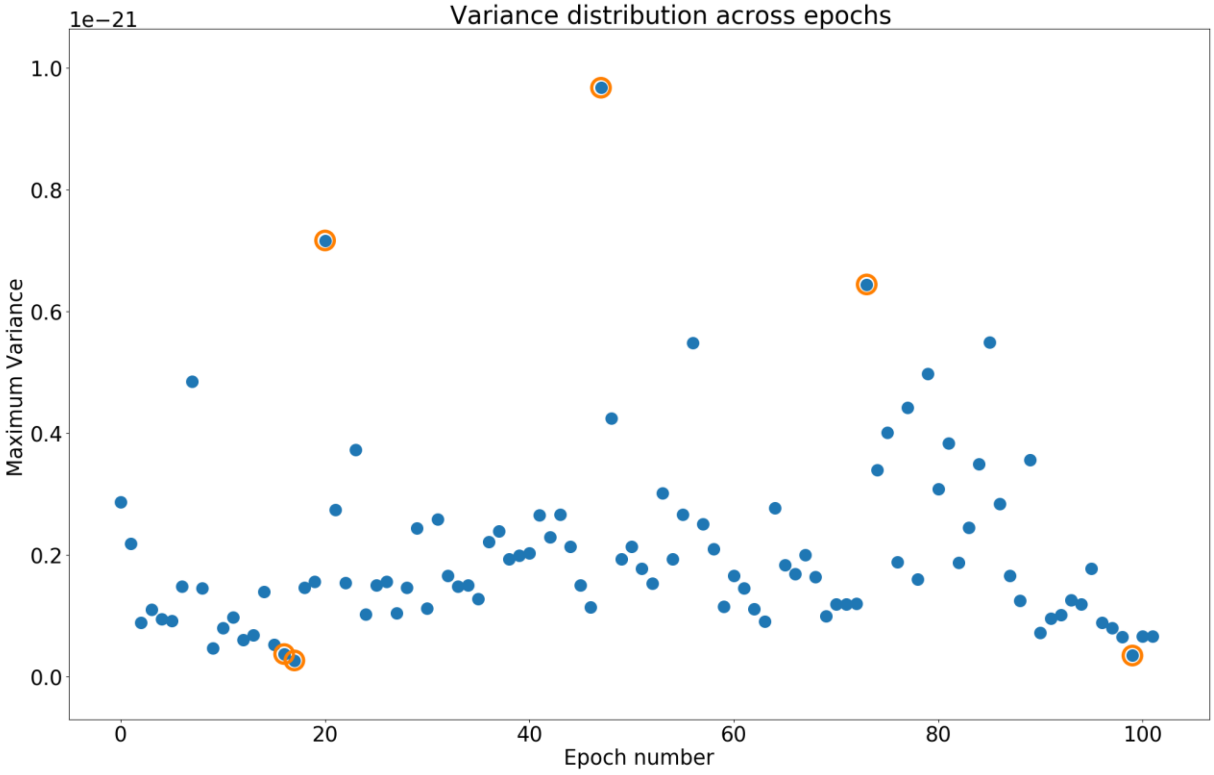


**Fig. 4**. Trial removal based on maximum covariance calculated over sensor array for all epochs, the red circled epochs are marked as bad and rejected from the study.

### 5. Changing sensor data unit from T and T/m to fT and fT/mm

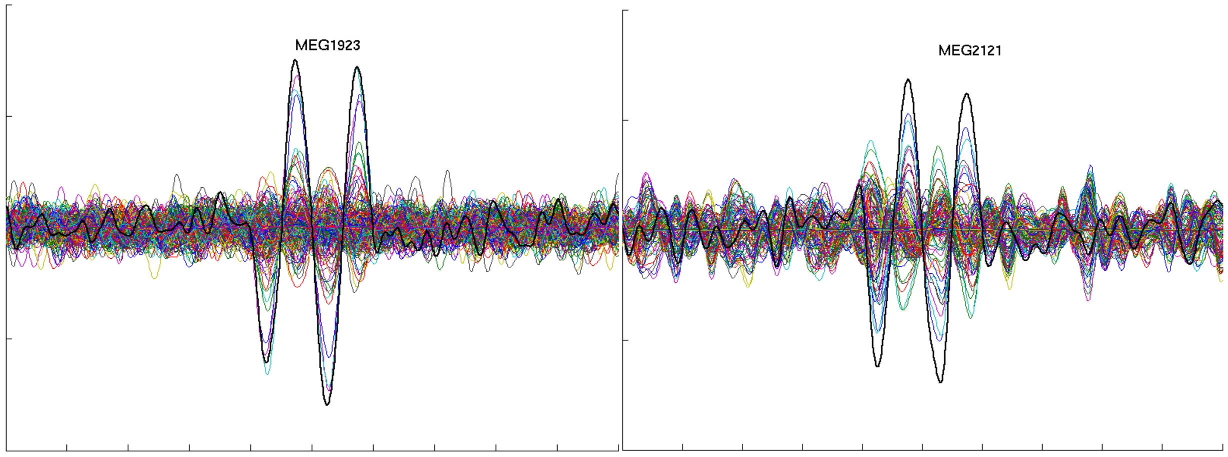


**Fig. 5.** Data conversion from T and T/m to fT and fT/mm turns the numeric value of magnetometer data higher.

### 6. Effect of filtering on data in different packages

<table>
<tr><td>MEG 2531</td><td>Baseline corrected raw data</td><td>Bandpass filtered 2-40Hz (default filter)</td></tr>
<tr><td>Without MaxFilte(R)</td><td>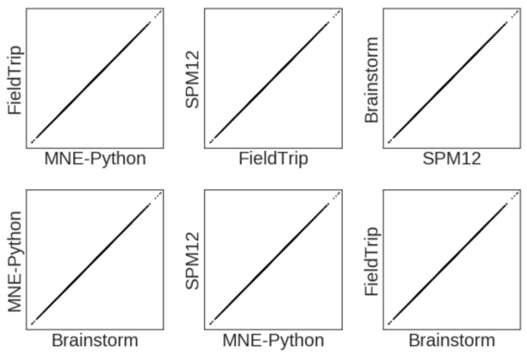
</td><td>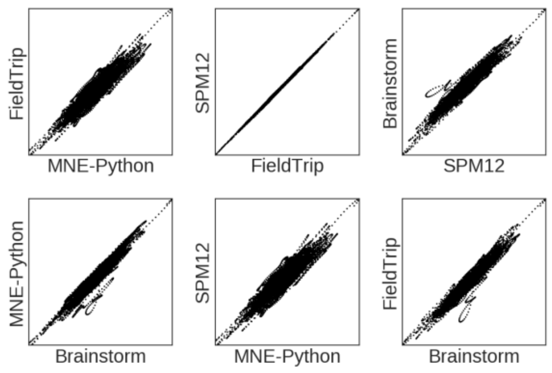
</td></tr>
<tr><td>After MaxFilte(R)</td><td>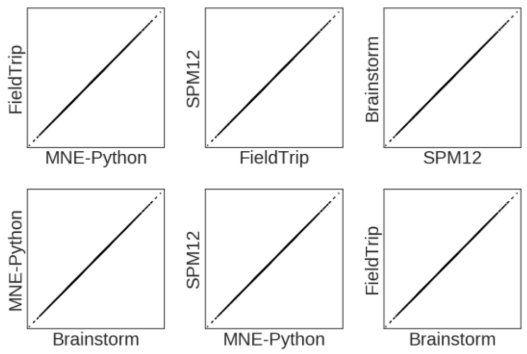
</td><td>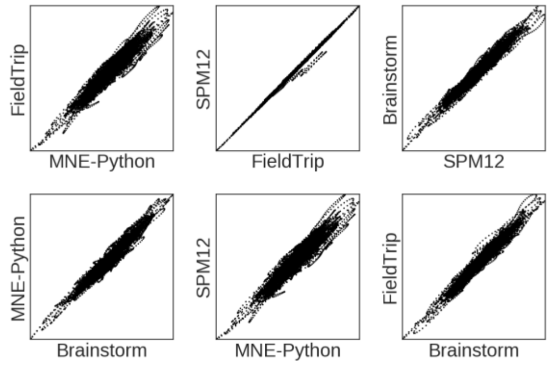
</td></tr>
</table>

(a)

<table>
<tr><td>MEG 1632</td><td>Baseline corrected raw data</td><td>Bandpass filtered 2-40Hz (default filter)</td></tr>
<tr><td>Without MaxFilte(R)</td><td>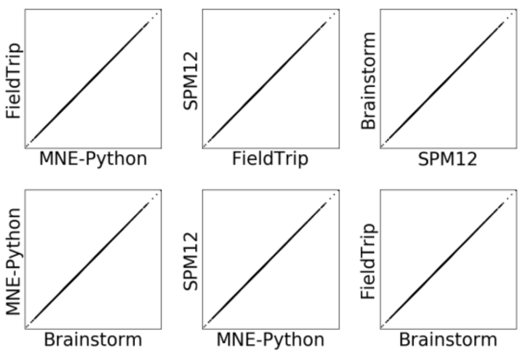
</td><td>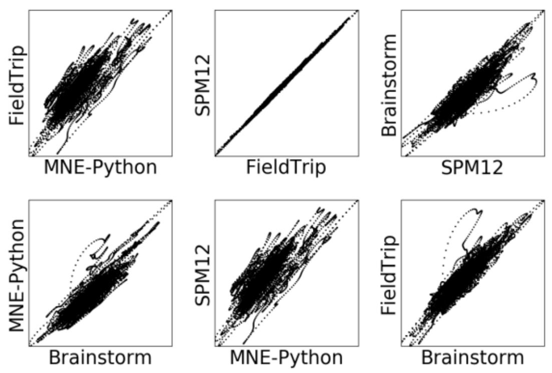
</td></tr>
</table>

| | | |
|---|---|---|
| After MaxFilte(R) | 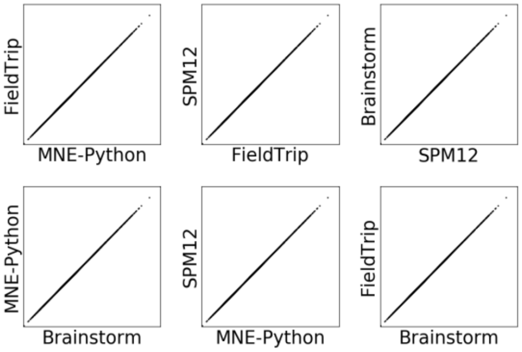  | 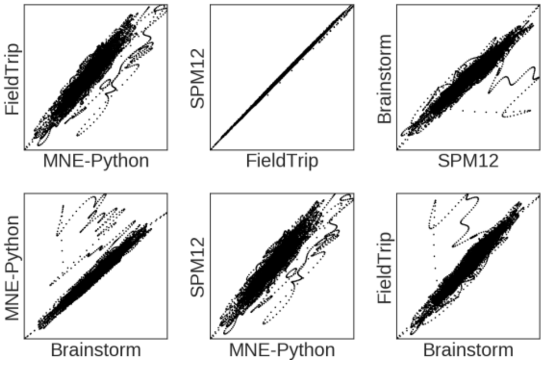  |

(b)

| MEG 0733 | Baseline corrected raw data | Bandpass filtered 2-40Hz (default filter) |
|---|---|---|
| Without MaxFilte(R) | 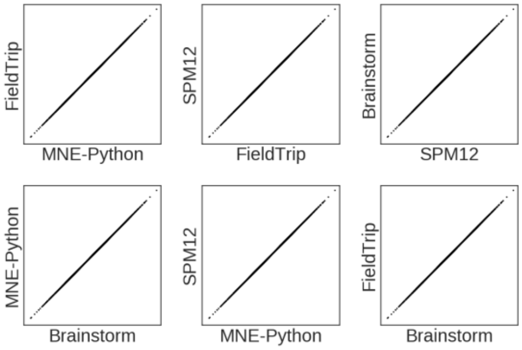  | 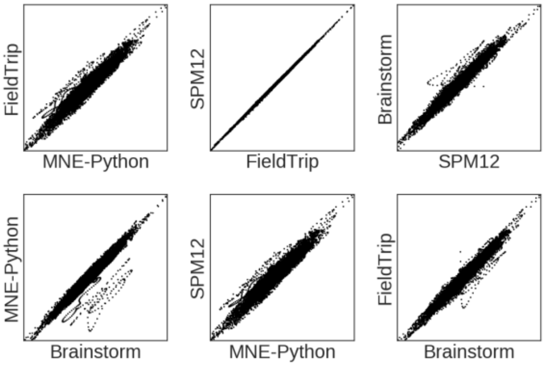  |
| After MaxFilte(R) | 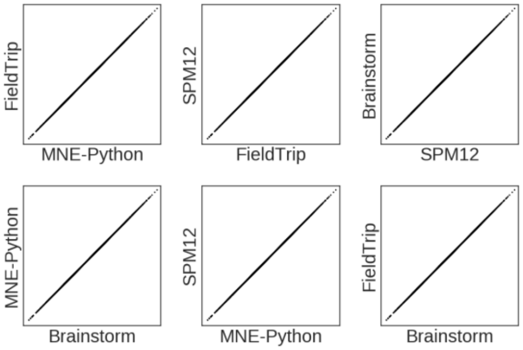  | 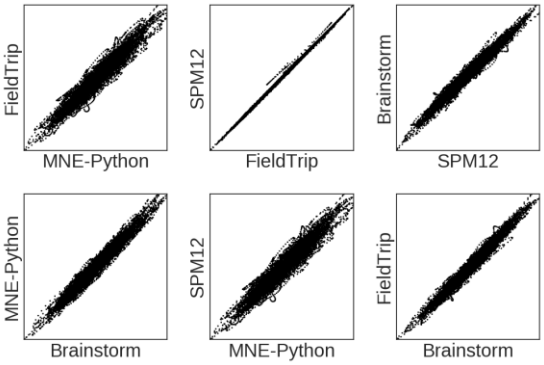  |

(c)

**Fig. 6.** Correlation between same input data across the four LCMV implementations, for randomly selected three channels. (a) Magnetometer (MEG2531); (b) Gradiometer (MEG1632); (c) Gradiometer (MEG0733)

### 7. Effect of SSS preprocessing on the rank estimation of the data

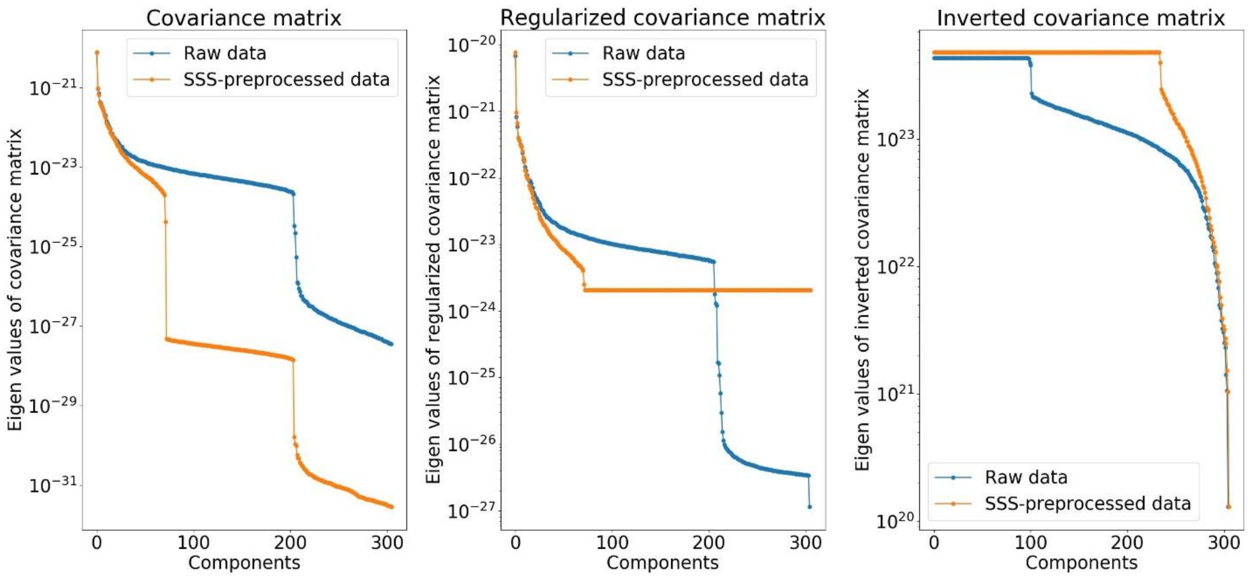


**Fig. 7**. Comparison of eigen spectrum for covariance matrix, regularized covariance matrix (with 5% of mean of trace) and inverted covariance matrix for raw and SSS-preprocessed data.

### 8. General workflow for beamformer analysis across the packages

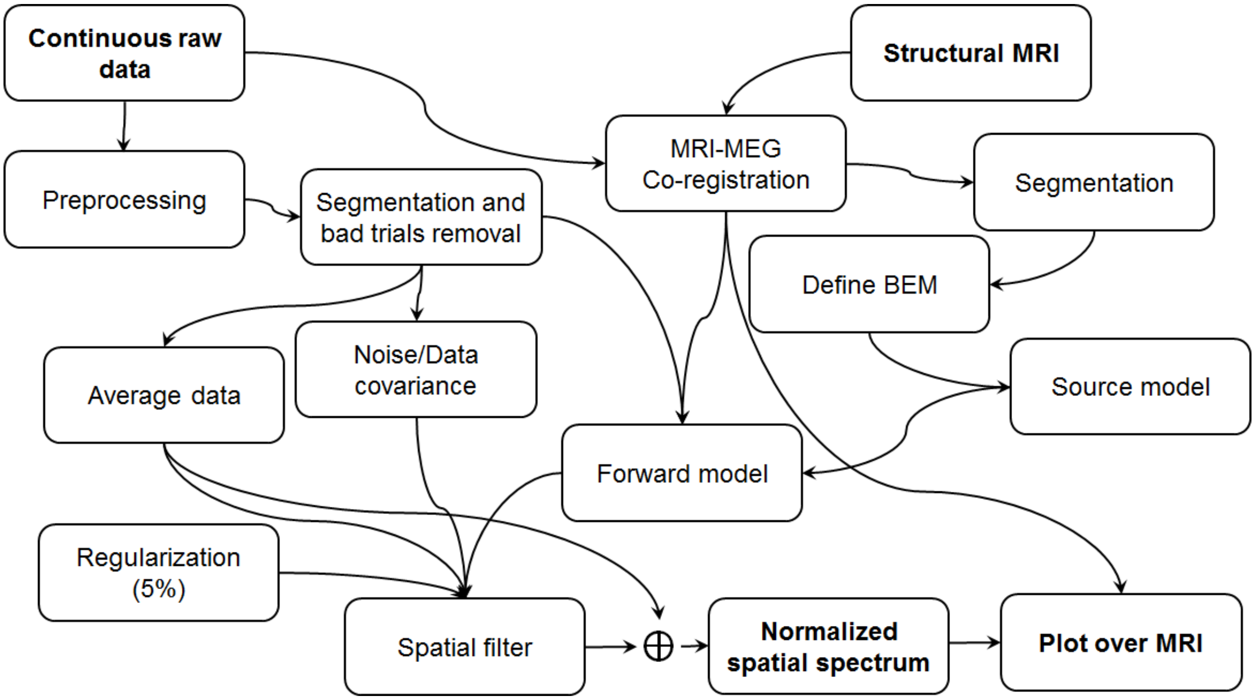


**Fig. 8**. Detailed general pipeline for LCMV beamformer for MEG/EEG source estimation used in all four packages

### 9. Point spread volume (PSV) for multimodal evoked field dataset estimated across the four LCMV implementations

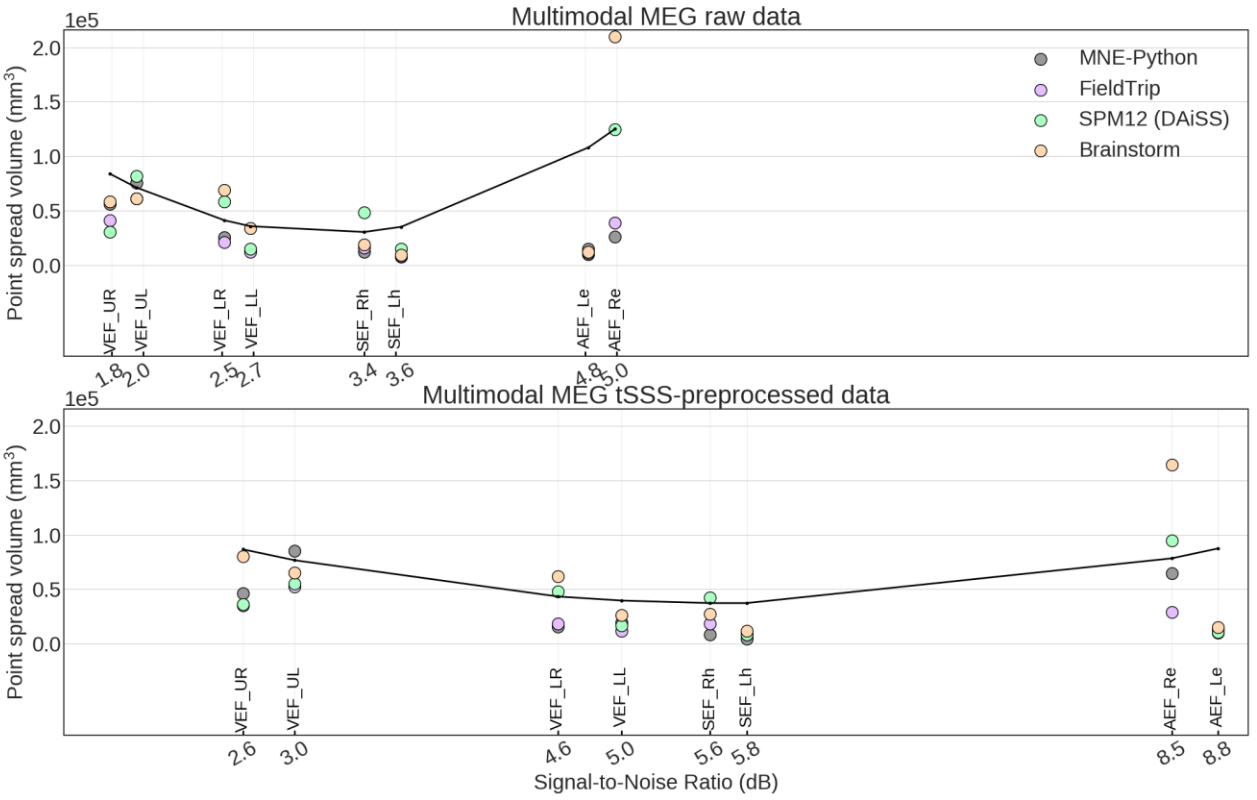


**Fig. 9**. PSV values estimated across the four LCMV implementations for multimodal evoked field raw data and tSSS preprocessed data with movement compensation.
